# Omics-assisted systematic exploration of the intricate regulatory network of guvermectin biosynthesis centered by the cluster-situated regulator GvmR in *Streptomyces*

**DOI:** 10.1101/2023.11.19.566996

**Authors:** Haoran Shi, Jiabin Wang, Shanshan Li, Chongxi Liu, Zhuoxu Dong, Xiangjing Wang, Yanyan Zhang, Wensheng Xiang

## Abstract

Guvermectin, produced by *Streptomyces* bacteria, is a purine nucleoside natural product recently registered as a new biopesticide to boost rice yield. Despite its importance, the regulatory network governing guvermectin biosynthesis remains largely unknown, severely impeding industrial-scale production and widespread application in rice production. Here, we investigated the diverse regulatory mechanisms employed by the cluster-situated regulatory gene, *gvmR*, in controlling guvermectin production from the perspective of widespread disturbance of gene expression at genome scale. GvmR activates the expression of guvermectin cluster by binding to the *gvmR*, *gvmA* and *O1* promoters. Additionally, GvmR binds to the promoter of *scnR1*, a previously unidentified and highly conserved regulator in *Streptomyces*. *scnR1* overexpression significantly suppressed guvermectin production by regulating the guvermectin cluster through binding to the same promoters as GvmR. Transcriptomic analysis revealed that GvmR extensively influences the expression of numerous genes located outside the guvermectin cluster, including the precursor supply (purine biosynthesis) and energy synthesis (oxidative phosphorylation) pathway genes, as well as 252 transcriptional regulatory genes. By genetic screening from 48 of these 252 regulatory genes, we identified additional five highly conserved genes that impact guvermectin production, suggesting a functional interplay between GvmR and highly conserved regulators in coordinating guvermectin production. These findings enrich our knowledge of the regulatory network governing guvermectin biosynthesis and offer a broadly applicable approach for investigating the molecular regulation of natural product biosynthesis and their high-titer production.

## INTRODUCTION

Guvermectin, a purine nucleoside natural product, is produced by several *Streptomyces* including *Streptomyces angustmyceticus*, *Streptomyces decoyicus* and *Streptomyces caniferus*, and possesses potent plant growth-promoting activities (Liu et al., 2022; Sugimori and Suhadolnik, 1965; Yuntsen et al., 1956; Yuntsen et al., 1954). A single seed-soaking treatment on 28 rice varieties with guvermectin can promote the growth of rice roots and coleoptile, tillering, and shorten the growth maturation, and ultimately boosts rice yield by 6.2% to 19.6%. This effect surpasses the yield achieved by other commercial growth-promoting biopesticides, such as cytokinin, brassinosteroid and the mixtures of gibberellin-auxin-brassinosteroid (Liu et al., 2022). Given its superior impact on rice growth and yield, guvermectin was registered as a new biochemical pesticide in China in 2021 (registration number PD20212929). Undoubtedly, the application of guvermectin presents new opportunities to enhance global crop yields.

In the past three years, the biosynthetic gene cluster (BGC) responsible for guvermectin production has been identified in *S. angustmyceticus* NBRC 3934, *S. decoyicus* NRRL 2666 and *S. caniferus* NEAU6, respectively (Liu et al., 2023; Shiraishi et al., 2021; Yu et al., 2021). The guvermectin BGCs in these strains share high similarity and comprise nine genes, including six biosynthetic key enzyme genes (*gvmA*, *gvmB*, *gvmC*, *gvmD*, *gvmE* and *gvmF*), one regulatory gene (*gvmR*), and two MFS (major facilitator superfamily) transporter genes (*gvmT1* and *gvmT2*). Comprehensive genetic and biochemical studies have been conducted to elucidate the guvermectin biosynthetic pathway, which proceeds via multiple steps utilizing D-fructose 6-phosphate and ATP as precursors (Liu et al., 2023; Yu et al., 2021). Initially, a D-psicose 6-phosphate 3-epimerase GvmD catalyzes the epimerization of D-fructose 6-phosphate to generate D-psicose 6-phosphate. Subsequently, the enzyme D-psicose 6-phosphate pyrophosphokinase GvmC utilizes D-psicose 6-phosphate and ATP to generate 6-phosphopsicosyl-2-pyrophosphate (PPPP) and AMP. AMP can then undergo hydrolysis by the AMP phosphoribohydrolase AgmA, producing the direct precursor adenine for the purine moiety of guvermectin. PPPP and adenine are then catalyzed by the adenine phosphoribosyltransferase GvmE to produce a key intermediate psicofuranine 6’-phosphate (PMP). Finally, PMP is successively catalyzed by the phosphatase GvmB and the dehydratase GvmF to produce the final product guvermectin. The MFS transporters GvmT1 and GvmT2 are speculated to facilitate the export of guvermectin from the cytoplasm into the medium. While the inactivation of either *gvmT1* or *gvmT2* leads to a significant decrease in guvermectin production, the specific functions of these two transporter genes remain to be determined (Liu et al., 2023). *gvmR* inactivation results in almost no production of guvermectin, suggesting the essential role of *gvmR* in guvermectin biosynthesis (Liu et al., 2023). However, the corresponding regulatory mechanism has not been elucidated.

Streptomycetes are a rich source for pharmaceutically valuable natural products (NPs). However, the initial titers of NPs are often low due to the stringent control exerted by intricate intracellular regulatory networks., DNA-binding transcriptional regulators, a pivotal component of these networks, play essential roles in determining NP production and final titers (Liu et al., 2013). Cluster-situated regulators (CSRs) are instrumental in regulating the expression of BGCs, thereby influencing the production level of the corresponding NPs (Liu et al., 2013). Typically, signal transduction pathways, predominantly orchestrated by global or pleiotropic regulators whose encoding genes are located outside the BGCs, moderate BGC expression by converging on the promoter regions of CSRs. Consequently, CSRs serve as direct and efficient switches for the expression of specific BGCs. Unveiling the regulatory mechanisms of CSRs not only facilitate the discovery and comprehension of new phenomena and action principles in living organisms, but more importantly, provides beneficial targets that can be engineered to improve NP production. The CSR-type regulator-based genetic engineering strategies have been extensively employed in the discovery and overproduction of NPs (Bu et al., 2021; Li and Tan, 2017). To date, multiple advances have been made in understanding of the novel and diverse regulatory mechanisms employed by CSRs to control the expression of cognate BGCs (e.g., cascade, feed-forward/feedback, and synergistic/coordinative effect) (Li et al., 2022; Li et al., 2023; Liu et al., 2013; Zhang and Tan, 2023). Many CSRs, such as *redZ* from the *S. coelicolor* Red cluster, *jadR1* from the *S. venezuelae* Jad cluster and *nemR* from the *S. cyaneogriseus* nemadectin cluster, exhibit pleiotropic roles affecting other NP BGCs, primary metabolism and even morphological development (Huang et al., 2005; Li et al., 2019; Xu et al., 2010). It is noteworthy that existing studies have primarily focused on the impact of CSRs on strain phenotype and the expression of a limited number of genes related to phenotype. Little is known about how CSR extensively alter gene expression at the genome level, and further exploration of targets beneficial for NPs production from CSR-disturbed gene expression changes is still a blank, which are imperative for a systematic understanding of the efficient biosynthesis of NPs and construction of high-producing industrial strains.

In this work, we employed omics analysis and overexpression screens to systematically elucidate the regulatory mechanisms of GvmR in guvermectin production. As a CSR, GvmR is not only essential for the expression of the guvermectin cluster, but also can bind directly to the promoter region of *scnR1*, a highly conserved regulatory gene belonging to the LacI family in *Streptomyces*. Intriguingly, overexpression of *scnR1* strongly inhibited guvermectin production. Moreover, transcriptomic analyses unveiled that the inactivation of *gvmR* has a profound impact on the expression of numerous genes situated outside of the guvermectin cluster. This includes genes involved in the precursor (purine biosynthesis) and energy synthesis (oxidative phosphorylation) pathways, as well as many transcriptional regulatory genes. Notably, some of these transcriptional regulatory genes were found to exert positive or negative regulatory effects on guvermectin production. These findings provide a partial yet insightful glimpse into the intricate regulatory network governing guvermectin production, with GvmR at its core. The identified regulatory interactions offer novel new avenues for exploring the underlying mechanisms and uncovering potential regulators for achieving high NP titers.

## RESULTS

### GvmR activates the expression of guvermectin BGC

*gvmR* is located upstream of, and in the opposite orientation as, the operon covering the six essential biosynthetic enzyme genes (*gvmA–F*) of guvermectin biosynthesis (Figure 1A) and encodes a LacI family regulatory protein consisting of 339 amino acids. GvmR contains a N-terminal LacI-type HTH motif and a C-terminal sensor (Peripla_BP_3) domain, which can respond to small molecules such as sugars, sugar phosphates, sugar acids and purines (Matilla et al., 2022). It was previously reported that *gvmR* inactivation caused substantial decrease of guvermectin production (Liu et al., 2023). To further verify that the dramatic titer decrease observed in the *gvmR* inactivation mutant (ΔgvmR) was indeed attributable to the absence of *gvmR*, a native copy of *gvmR* and another copy of *gvmR* driven by the constitutive *hrdB* promoter were inserted into the integrative plasmid pIJ10500 to generate pIJ10500::gvmR and pIJ10500::P_hrdB_gvmR, respectively. These two constructs were further introduced into ΔgvmR and the empty vector pIJ10500 was introduced into NEAU6 to obtain complementary strains ΔgvmR/gvmR and ΔgvmR/P_hrdB_gvmR and the control strain NEAU6/pIJ10500 (Figure 1B). The two complementary strains, two control strains (NEAU6 and NEAU6/pIJ10500) and ΔgvmR were cultured in fermentation medium for 8 days, and guvermectin production was quantified by HPLC analysis. Consistent with previous findings, *gvmR* inactivation resulted in the production of only trace amounts of guvermectin. Conversely, complementation of *gvmR* in ΔgvmR led to a substantial increase in guvermectin, reaching levels of hundreds of milligrams per liter (Figure 1C). The introduction of pIJ10500, in fact, exhibited an adverse effect on guvermectin production. Compared with the parental strain NEAU6, the guvermectin titer decreased by 30% in NEAU6/pIJ10500 (Figure 1C). Therefore, in comparison with the accurate control strain NEAU6/pIJ10500 (441 mg L^-1^), ΔgvmR/gvmR and ΔgvmR/P_hrdB_gvmR restored guvermectin production by 60% (263 mg L^-1^) and 105% (461 mg L^-1^), respectively (Figure 1C). These data indicated that GvmR plays a positive role in guvermectin production.

**Figure 1.**
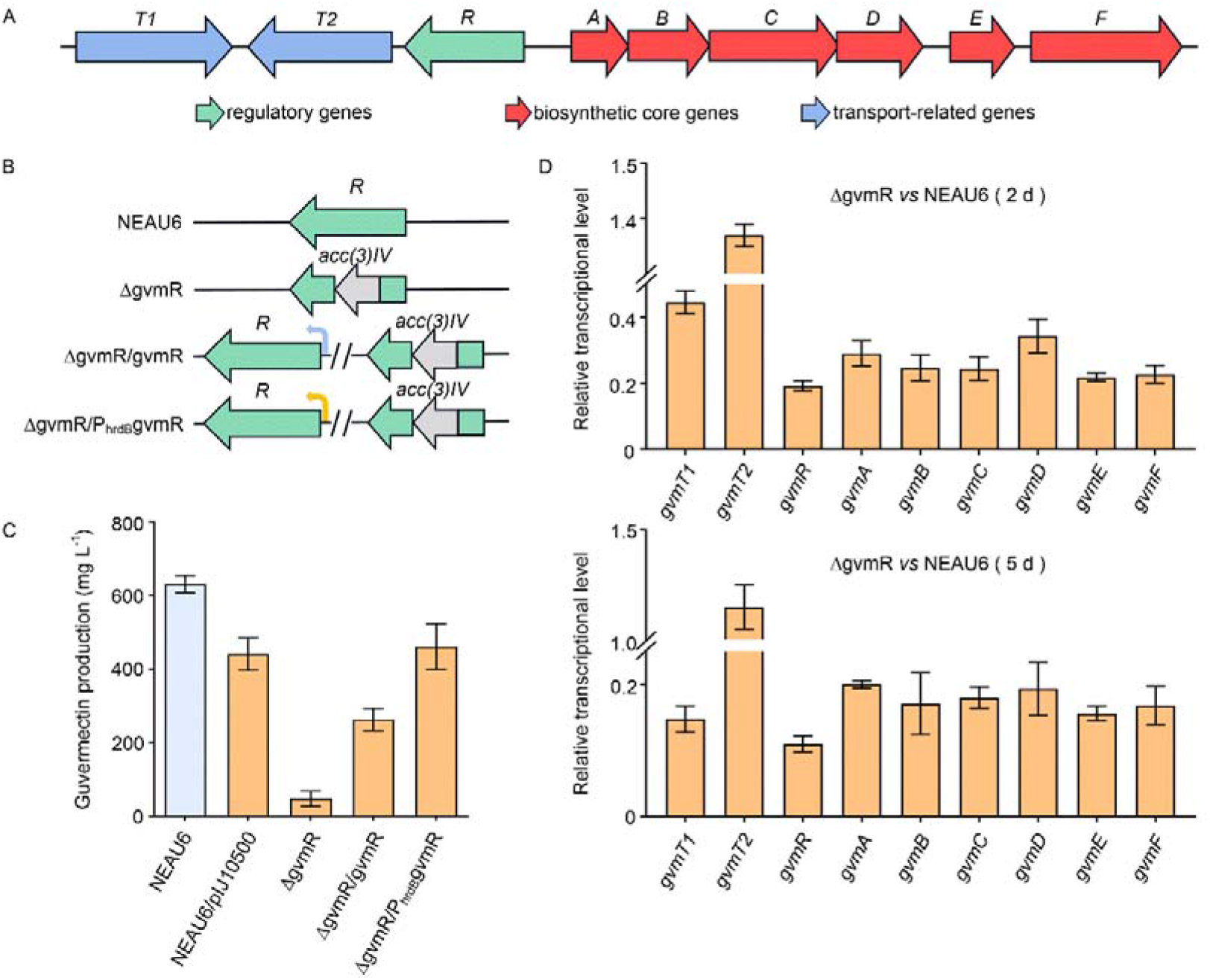
Effect of *gvmR* on the production of guvermectin and the expression of guvermectin BGC. A, genetic organization of guvermectin biosynthetic gene cluster. Each arrow indicates a separate open reading frame (ORF) and orientation of transcription. B, diagram of *gvmR* inactivation and complementation constructions. C, comparative guvermectin production in NEAU6, NEAU6/pIJ10500, ΔgvmR and two complementation strains (ΔgvmR/gvmR and ΔgvmR/P_hrdB_gvmR). D, qRT-PCR analysis determining the effects of *gvmR* on the transcription of guvermectin BGC genes. RNA samples were isolated from 2-and 5-days cultures. Transcription of each gene was calculated relative to the NEAU6 value on day 2 or day 5, assigned a value of 1. Transcription of 16SrRNA was used as the internal control.

As a CSR, GvmR may control guvermectin production by modulating the expression of some guvermectin BGC genes. To determine the effect of GvmR on expression of guvermectin biosynthesis genes, total RNAs were isolated from the mycelia of NEAU6 and ΔgvmR cultured for 2 and 5 days, and qRT-PCR was performed to assess expression of relevant genes. The results showed that transcript levels of six structural genes (*gvmA* to *gvmF*), *gvmR* and one predicted exporter gene *gvmT1* were all significantly lower in ΔgvmR than in NEAU6 at both time points (Figure 1D). On the contrary, the transcript level of another exporter gene, *gvmT2*, exhibited a slight elevation in ΔgvmR than in NEAU6 (Figure 1D), which might be explained by the insertional inactivation of *gvmR* ORF by the apramycin resistance gene *acc3* (Figure 1B). Thus, these data demonstrated that GvmR positively controls guvermectin production by activating the expression of guvermectin BGC genes.

### GvmR binds to the bidirectional *gvmR****–****A* and *O1* promoters

Identifying the direct targets of GvmR is important to understand how it activates the expression of guvermectin BGC genes. To this end, a thorough analysis of putative operons and promoter regions within the guvermectin BGC was conducted. In the cluster, the six key structural genes (*gvmA* to *gvmF*) are transcribed in the same direction. Notably, and the spacer lengths between *gvmD* and *gvmE*, as well as *gvmE* and *gvmF*, exceed 100-bp (Figure 2A), so we speculated that these six genes may constitute a polycistronic unit, though *gvmE* and *gvmF* might also possess their own promoters. *gvmT2* and *gvmR* are transcribed in the same direction, with a 61-bp between them, suggesting potential co-transcription. However, the possibility of *gvmT2* having its native promoter cannot be ruled out. Concerning *gvmT1*, located at the left end of the guvermectin BGC, an examination of the NEAU6 genome sequence revealed three convergently transcribed genes upstream of *gvmT1*: *scn1134* (*O1*, encoding a nicotinamidase-related amidase), *scn1135* (*O2*, encoding arginase), and *scn1136* (*O3*, encoding nitroreductase) (Figure 2A). The space length between *gvmT1* and the adjacent *O1* is 126-bp (Figure 2A). This suggests the possibility of *gvmT1* having independent promoter activity, but it could also be controlled by the *O1* promoter. To confirm our speculations and determine the authentic operons and promoter regions, RT-PCR-based co-transcription analysis and β-glucuronidase-based GUS assays were carried out. For co-transcriptional analysis, six primer pairs capable of amplifying the intergenic regions (regions 1 to 6) of adjacently located genes were designed (Figure 2A and Table S1). Then RT-PCR was performed using the RNAs prepared from the mycelia of NEAU6 fermented for 2 days. As anticipated, PCR products spanning the intergenic regions of *O1*/*O2*, *O3*/*gvmT1*, *gvmT2/gvmR*, *gvmD/gvmE* and *gvmE*/*gvmF* were detected (Figure 2B), indicating that the nine guvermectin BGC genes, i.e., *gvmT1*, *gvmR* to *gvmT2*, and *gvmA* to *gvmF* are organized into three polycistronic units: *O1*–*gvmT1*, *gvmR*–*gvmT2* and *gvmA*–*gvmF* (Figure 2A). For GUS assays, the upstream regions of four genes (*gvmT1*, *gvmT2*, *gvmE* and *gvmF*) were separately cloned upstream of the *gusA* gene (encoding β-glucuronidase) based on the integrative pSET152. Subsequently, these constructs along with the control plasmid pSET152::gusA (containing only the *gusA* ORF), were introduced into *Streptomyces coelicolor* M1146, respectively, to detect the corresponding β-glucuronidase activities. Compared with the control strain (harboring pSET152::gusA) that exhibited no GUS activity, the upstream regions of *gvmT1*(P_T1_) and *gvmF* (P_F_) gave strong GUS activities. the upstream regions of *gvmT2* (P_T1_) and *gvmE* (P_E_) also gave relatively weak GUS activities (Figured 2C). These results indicate that each of the four genes is regulated not only by its corresponding upstream polycistronic promoters but also by its individual promoters.

**Figure 2.**
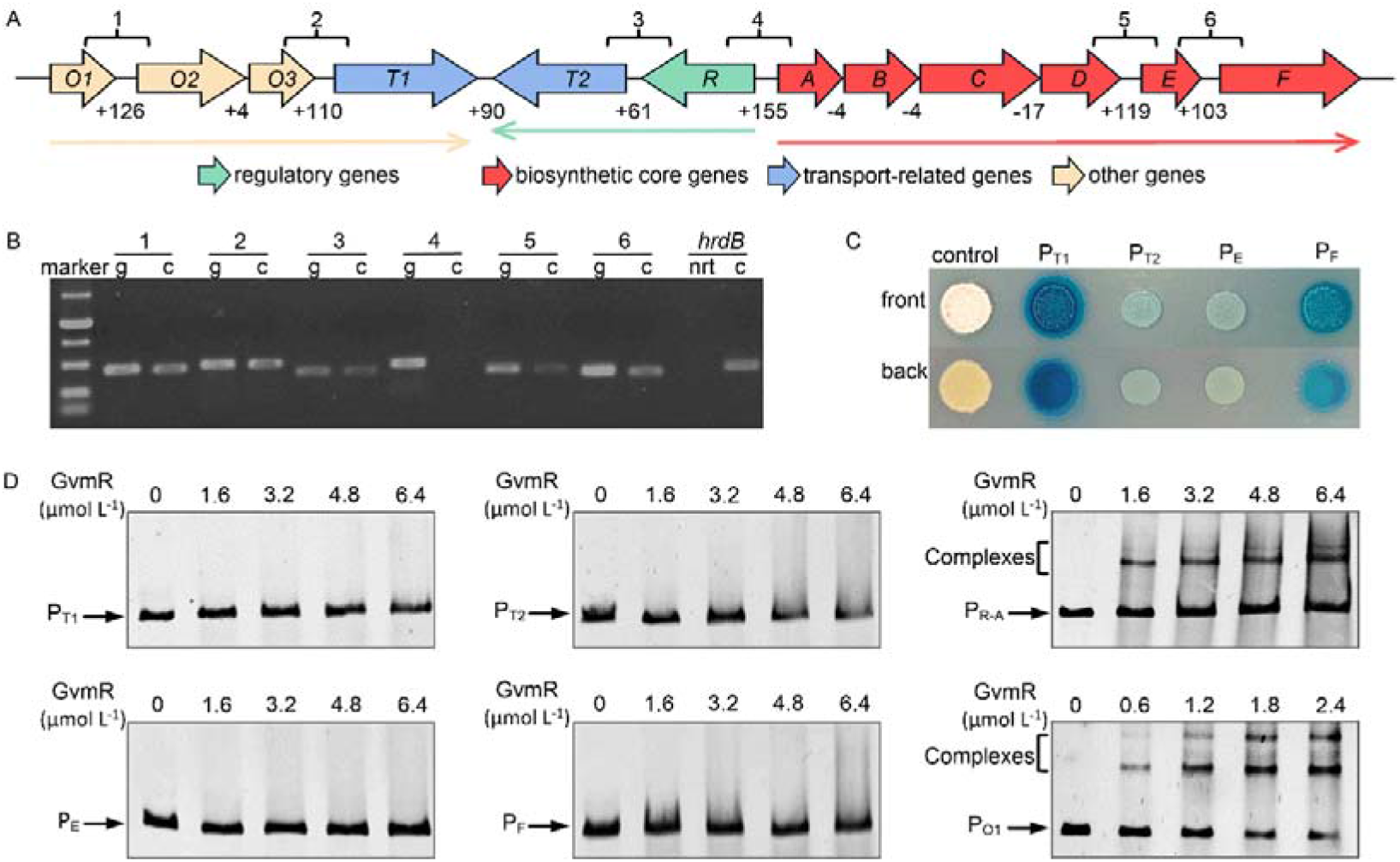
EMSAs to determine the direct targets of GvmR. A, the extended guvermectin BGC added with three genes possibly co-transcribed with *gvmT1*. The intergenic regions labelled 1–6 were used for RT-PCR amplification; three operons are represented by horizontal long arrows of different colors; the numbers below the guvermectin BGC indicate the space length (bp) of adjacent genes. B, co-transcription analysis of the guvermectin BGC. cDNA was generated from the total RNAs of NEAU6 strain cultured for 2 days. g, genomic DNA. c, cDNA template. nrt, “cDNA” template generated without reverse transcriptase. C, determination of the native promoters of *gvmT1*, *gvmT2*, *gvmE* and *gvmF* by GUS assays. D, EMSAs of the interactions between GvmR and the guvermectin BGC promoter regions. Arrows, free DNA probes. Square brackets, GvmR-DNA complexes.

Based on the findings from co-transcriptional analysis and β-glucuronidase assays, it is evident that the extended guvermectin cluster (encompassing *O1*, *O2* and *O3* as part of the guvermectin cluster) contains at least seven promoter regions. These promoter regions were then used for EMSAs with N-terminal GST-tagged GvmR to identify direct binding targets (Fiugre S1). Figure 2D illustrates that GvmR forms complexes with P_R-A_ (the bidirectional *gvmR*–*gvmA* promoter) and P_O1_ (the promoter region of *O1*). In contrast, no complex was observed for the other four promoter regions, indicating that GvmR directly activates transcription of the guvermectin cluster by binding to the promoter regions of *gvmR*, *gvmA* and *O1*.

### *In vitro* and *in vivo* confirmation of GvmR binding sequences

Usually, LacI/Gal family regulators interact with palindromic sequences within target promoter regions to exert control over gene expression (Schumacher et al., 1994; Swint-Kruse and Matthews, 2009). Based on this knowledge, we tried to search for potential palindromic sequences from P_R-A_ and P_O1_. As expected, four highly similar 14-bp palindromic sequences (sites I and II in P_R-A_; sites III and IV in P_O1_) were detected from the two promoter regions (Figure 3A). The sequence of site I is the same as that of site II; site III, when compared with site I, exhibits two variations at position 7, where the base is “G” instead of “A”, and at position 7 [where the base is “C” instead of “T” (Figure 3B); site IV, differs from site III at position 2 (2’) (Figure 3B). To ascertain the indispensability of the 14-bp palindromic sequence for GvmR’s DNA binding activity, we selected P_R-A_ as a representative and conducted site mutation experiments, resulting in the generation of three mutated DNA probes: P_R-A_-M1, P_R-A_-M2 and P_R-A_-M3 (Figure 3C). In P_R-A_-M1, site I was replaced by NdeI restriction site, while in P_R-A_-M2, site II was substituted by NheI restriction site. Meanwhile, P_R-A_-M3 featured mutations at both site I and site II, being replaced by NdeI and NheI restriction sites, respectively. These mutated DNA probes, along with the wild-type P_R-A_, were then subjected to EMSAs with GvmR, As shown in Figure 3D, P_R-A_ formed a single shifted band with GvmR at lower protein concentrations, and at a higher GvmR concentration (2.4 µM), a new, faintly shifted band appeared atop the original one. In contrast to P_R-A_, both P_R-A_-M1 and P_R-A_-M2 form only one band with GvmR, and P_R-A_-M3 completely lost its ability to form complexes with GvmR. This observation strongly suggests that the two palindromic sequences are indispensable for P_R-A_ binding with GvmR. Subsequently, to evaluate the necessity of these 14-bp palindromic sites for P_R-A_ promoter activity and further validate their importance in mediating GvmR’s regulatory effects on *gvmA*-oriented gene expression *in vivo*, we designed a truncated P_R-A_ promoter probe, P_A_, which was 142-bp shorter than P_R-A_ (Figure 4A). Three mutants of P_A_, denoted as P_A_-M1, P_A_-M2 and P_A_-M3, derived from P_R-A_-M1, P_R-A_-M2, and P_R-A_-M3, respectively, were also prepared. P_A_ and the three mutant promoters, were inserted separately upstream of *gusA* in pSET152 to yield pSET152::P_A_gusA, pSET152::P_A_-M1gusA, pSET152::P_A_-M2gusA and pSET152::P_A_-M3gusA. Subsequently, *gvmR* controlled by P_hrdB_ was assembled into the above four recombinant plasmids to generate another four plasmids pSET152::P_A_gusA::P_hrdB_gvmR, pSET152::P_A_-M1gusA::P_hrdB_gvmR, pSET152::P_A_-M2gusA::P_hrdB_gvmR and pSET152::P_A_-M3gusA::P_hrdB_gvmR. These four constructs together with pSET152::P_A_gusA were introduced into ΔgvmR, generating strains ΔgvmR/P_A_gusA::P_hrdB_gvmR, ΔgvmR/P_A_-M1gusA::P_hrdB_gvmR, ΔgvmR/P_A_-M2gusA::P_hrdB_gvmR, ΔgvmR/P_A_-M3gusA::P_hrdB_gvmR, ΔgvmR/P_A_gusA, respectively. All strains obtained above were cultured in guvermectin fermentation medium and their GUS activities were quantitatively measured using the substrate p-nitrophenyl-β-D-glucuronide. In the absence of GvmR expression (ΔgvmR/P_A_gusA), P_A_ failed to activate the expression of *gusA*. Conversely, in the presence of constitutive GvmR expression (pSET152::P_A_gusA::P_hrdB_gvmR), transcription of *gusA* driven by P_A_ was readily detected (Figure 4B and 4C), confirming the direct activation of P_A_ by GvmR. Moreover, under conditions of GvmR constitutive expression, mutations at site I and II of P_A_ led to a significant decrease in GUS activities; simultaneous mutation of both sites resulted in a more severe reduction (Figure 4B and 4C). These results clearly indicated that the two 14-bp palindromic sites play a crucial role in mediating the activation of P_A_ by GvmR.

**Figure 3.**
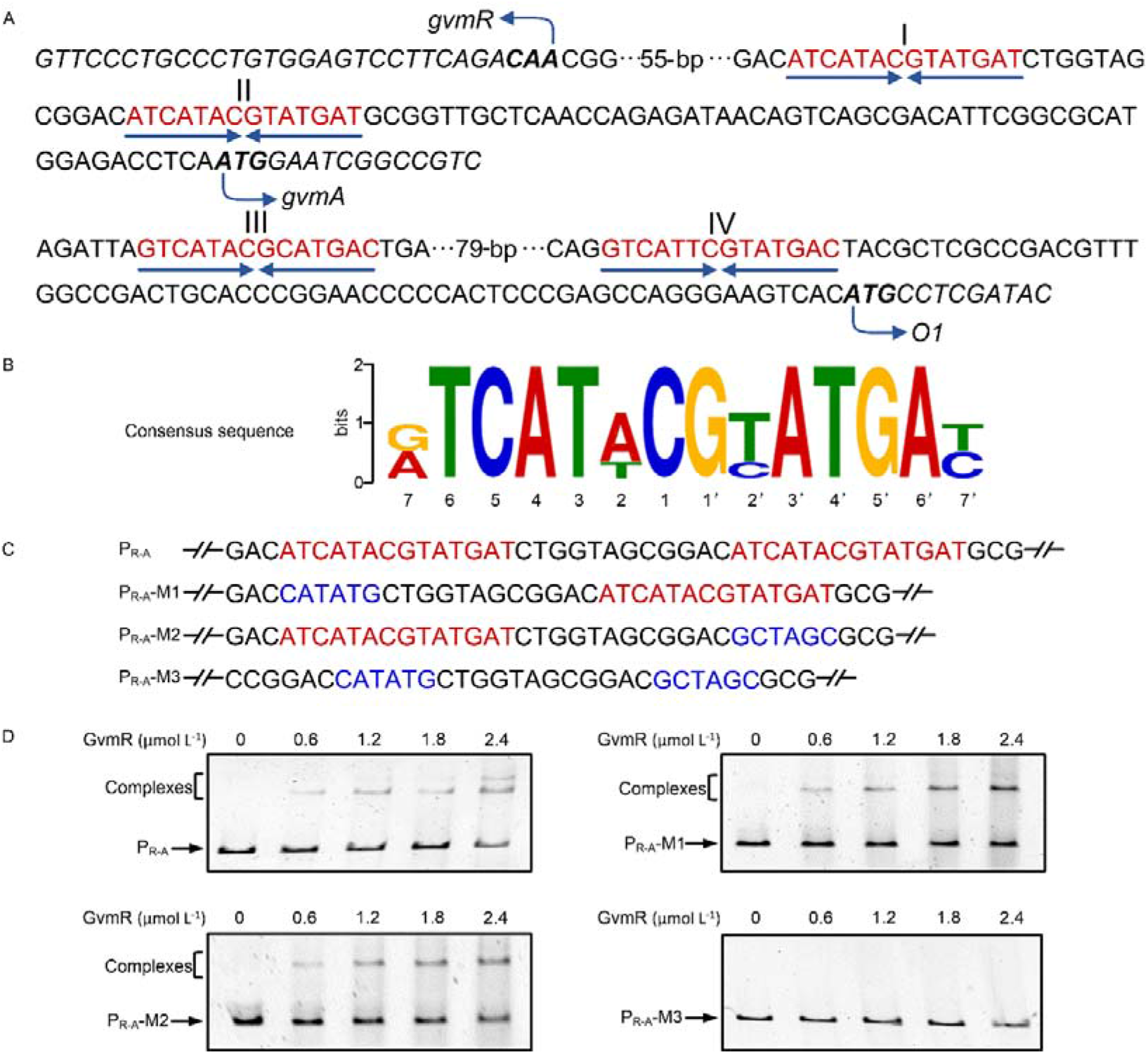
Determination of GvmR conservative binding sites. A, four similar 14-bp palindromic sequences discovered from the intergenic region of *gvmR*-*gvmA* (P_R-A_) and the upstream region of *O1* (P_O1_). The translational start sites of *gvmR*, *gvmA* and *scnO1* were indicated by bent arrows. The palindromic sequences are in red and marked by dark blue arrows. B, conservative sequence analysis of GvmR binding sites. C, mutation diagram of the two palindromic sequences in P_R-A_. The changed nucleotides are in blue. D, EMSAs of GvmR binding to P_R-A_ and the mutated probes P_R-A_-M1, P_R-A_-M2, and P_R-A_-M3.

**Figure 4.**
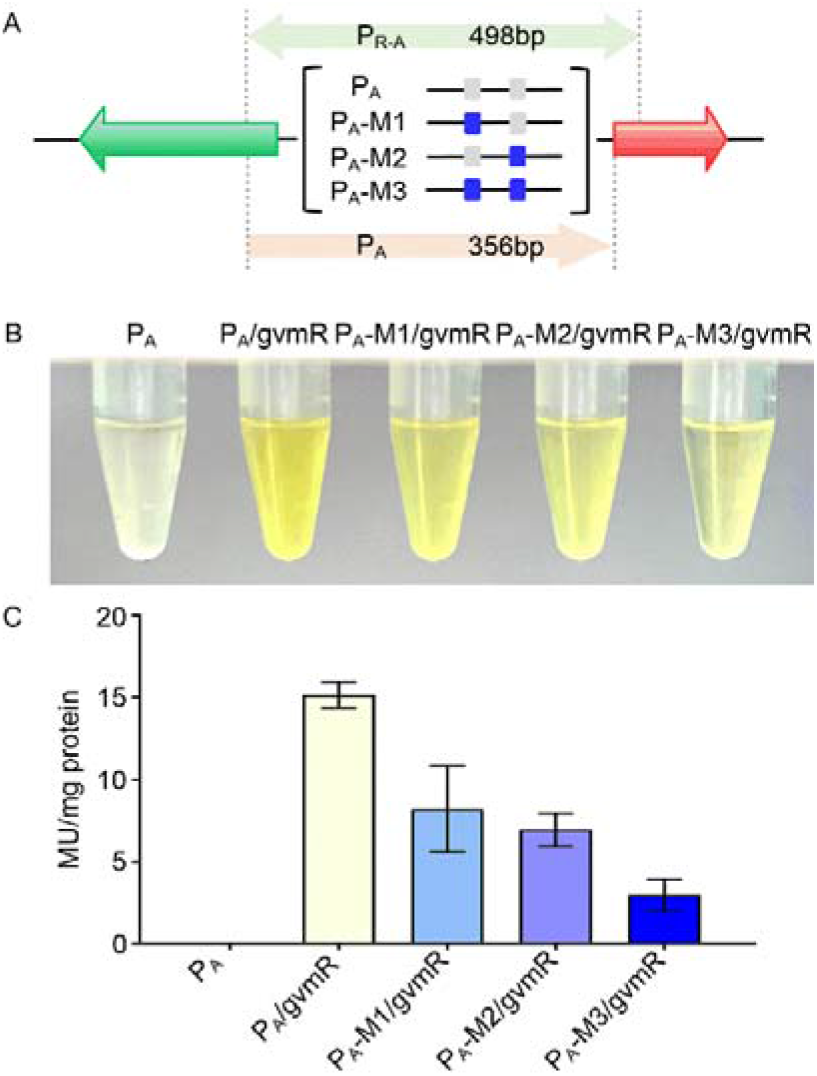
GUS assays to confirm the importance of the palindromic sequences for GvmR mediated activation of P_A_. A, mutation diagram of the two palindromic sequences in P_A_. Grey box indicates the wild-type palindromic sequences; blue box indicates the site replaced by NdeI or NheI restriction site. B, GUS assays to detect the effect of site mutation on GvmR mediated activation of P_A_ with p-nitrophenyl-β-D-glucuronide as the substrate. C, spectrophotometric assays of the effect of site mutation on GvmR’s activation of P_A_. P_A_: ΔgvmR/P_A_gusA; P_A_/gvmR: ΔgvmR/P_A_gusA::P_hrdB_gvmR; P_A_-M1/gvmR: ΔgvmR/P_A_-M1gusA::P_hrdB_gvmR; P_A_-M2/gvmR: ΔgvmR/P_A_-M2gusA::P_hrdB_gvmR; P_A_-M3/gvmR: ΔgvmR/P_A_-M3gusA::P_hrdB_gvmR.

### *scnR1*, a new highly conserved lacI-like regulator, is a direct target of GvmR, and its overexpression represses guvermectin production

As mentioned above, a 14-bp palindromic sequences were confirmed to be necessary for the DNA-binding activity of GvmR. We thus wonder that whether there are such similar palindromic sequences bound by GvmR in the promoter regions beyond the guvermectin cluster. To address this inquiry, the 14-bp palindromic sequences was used to scan the NEAU6 genome with MAST/MEME (http://meme-suite.org). This search yielded a total of 54 promoter regions that harbored intact or partial sequences resembling the 14-bp binding motifs (Table S2). These regions were subsequently subjected to EMSAs with GvmR, revealing interaction with only one specific promoter probe P_R1_, located in the promoter region of *scnR1* (*scn1544*) (Figure 5A and 5B). Remarkably, this *scnR1* promoter contained two 14-bp palindromic sequences separated by a 7-bp interval (Figure 5A). ScnR1, a LacI family transcriptional regulator, shows strong similarity to other LacI members, such as *sco1642* (68% identity) from *Streptomyces coelicolor* and *sbi_8484* (75% identity) from *Streptomyces bingchenggensis*. An intriguing observation was the high conservation of *scnR1* and its orthologs across *Streptomyces* species. Through a search encompassing 268 *Streptomyces* genomes obtained from the NCBI Genome database (as of Aug. 2021), we identified 263 ScnR1 orthologs (query coverage 90% to 100% and percent identity 60% to 100%) present in 261 *Streptomyces* species. Nevertheless, despite this conservation, their functional roles remain largely unexplored. qRT-PCR was then performed to determine the regulatory effect of GvmR on *scnR1* expression. However, *gvmR* inactivation did not result in a significant alteration in *scnR1* transcription (Figure 5C).

**Figure 5.**
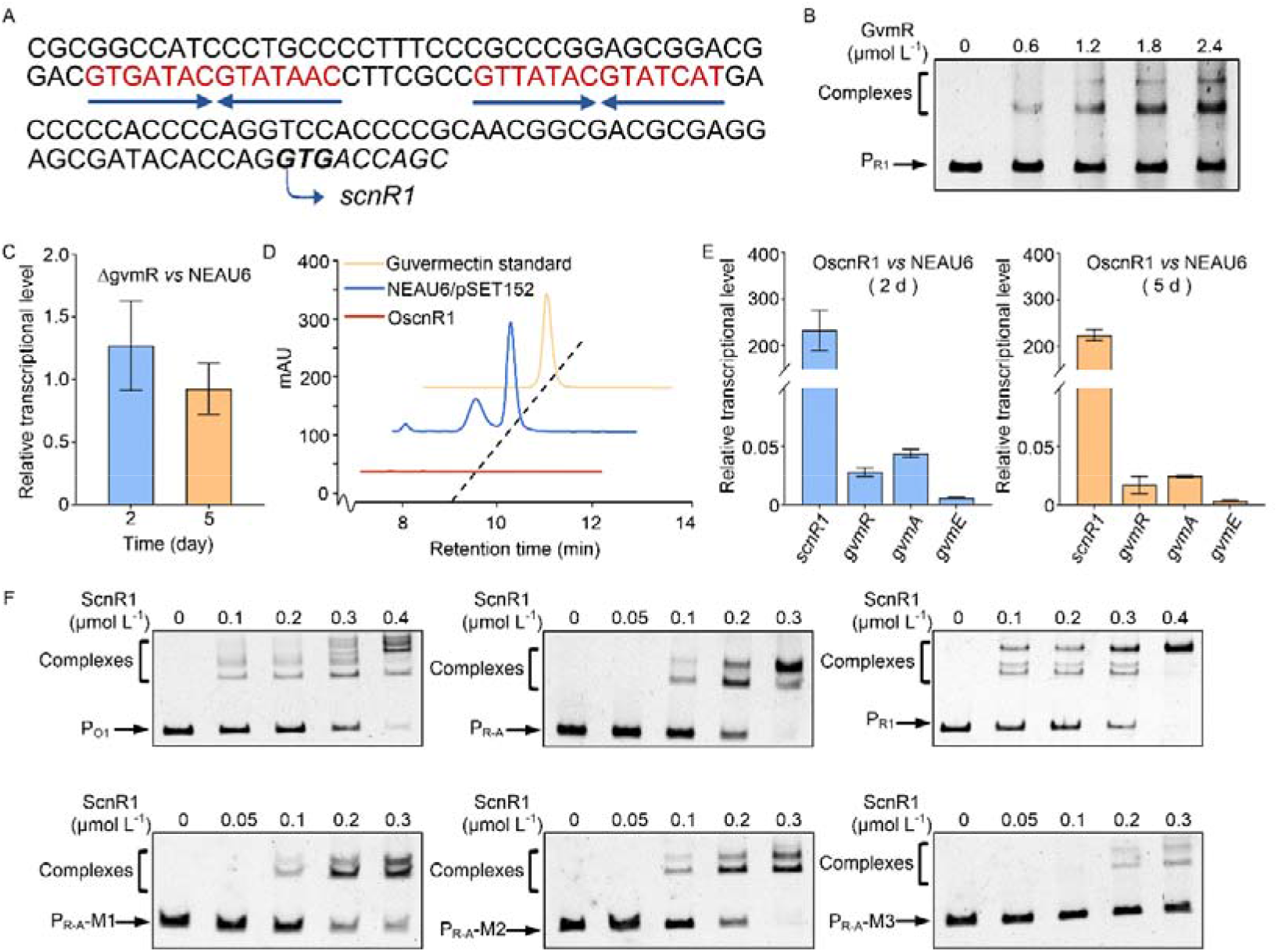
Effect of *scnR1* overexpression on the production of guvermectin and the expression of guvermectin BGC. A, characteristic sequence analysis of the promoter region of *scnR1*. The translation start sites of *scnR1* were indicated by bent arrows. The palindromic sequences are in red and marked by dark blue arrows. B, EMSAs of GvmR binding to the promoter region of *scnR1* (P_R1_). C, effect of *gvmR* inactivation on the transcription of *scnR1*. Transcription of *scnR1* was calculated relative to the NEAU6 value on day 2 or day 5, which was assigned a value of 1. 16SrRNA transcription was used as the internal reference. D, HPLC analysis of guvermectin production in the control strain NEAU6/pSET152 and *scnR1* overexpression strain OscnR1. E, transcriptional analysis of *scnR1*, *gvmR*, *gvmA* and *gvmE* in strains NEAU6 and OscnR1. The transcriptional level of each gene in NEAU6 at each time point was assigned as 1. F, EMSAs of ScnR1 binding to its targets and identification of its binding sites.

Considering ScnR1’s status as a highly conserved transcriptional regulator in *Streptomyces*, the question arises: could ScnR1 influence guvermectin production? To explore this prospect, a *scnR1* overexpression strain OscnR1, in which *scnR1* expression was controlled by the *hrdB* promoter in an integrative pSET152, was constructed. Indeed, after flask fermentation, no guvermectin was produced by OscnR1 in comparison with NEAU6 and NEAU6/pSET152 (Figure 5D), indicating that *scnR1* overexpression has strong repressive effect on guvermectin production.

### Identification of ScnR1 targets and its binding sequences

To elucidate the repressive mechanism underlying guvermectin biosynthesis resulting from the overexpression of *scnR1*, we conducted three experiments: qRT-PCR analysis of several genes (*gvmR*, *gvmA* and *gvmE*) to determine the influence of ScnR1 on guvermectin BGC, EMSAs to find the direct targets of ScnR1, and promoter mutational analysis to determine the characteristic binding sequences of ScnR1. The qRT-PCR results revealed a significant reduction in the transcript levels of all three genes in OscnR1 compared to NEAU6 at both time points (Figure 5E), indicating the inhibitory effect of ScnR1 on the expression of the guvermectin BGC. To ascertain whether this inhibition is direct or indirect, EMSAs were carried out with all the promoter regions within the guvermectin BGC using GST-tagged ScnR1. Figure 5F demonstrates that ScnR1 binds to the promoter regions of P_R-A_, P_O1_ and P_R1_ in a concentration-dependent manner. Notably, these three promoter probes have been previously as direct targets of GvmR (Figure 2), establishing that ScnR1 and GvmR can bind to the same target promoters. This observation promoted us to investigate if ScnR1 shares similar characteristic binding sequences with GvmR. To test this hypothesis, three mutated probes (P_R-A_-M1, P _R-A_-M2, and P_R-A_-M3) depicted in Figure 3 were individually subjected to EMSAs with ScnR1. Compared with the wild-type P_R-A_, all three mutated probes formed visible fewer DNA-ScnR1 complexes (Figure 5F); Notably, P_A_-M3 containing mutations at both site 1 and site 2, exhibited the weakest interactions with ScnR1, indicating that the two 14-bp palindromic sequences are essential for the binding activity of ScnR1.

### GvmR exerts significant influence on genes related to ribosome, oxidative phosphorylation and purine biosynthesis

The direct interaction between GvmR and the promoter region of the highly conserved *scnR1* suggests that GvmR may function as a pleiotropic regulator. To gain deeper insights into the pleiotropic functions of GvmR and to explore the intrinsic regulatory mechanisms underlying the inability of ΔgvmR to produce guvermectin, we conducted comparative RNA-seq analysis using total RNAs isolated from NEAU6 and ΔgvmR strains cultured for various days (0.5, 1, 2 and 5 days). Statistical analysis revealed that, employing criteria of |log_2_ (fold change) | > 0.5 and *p*_adj_ < 0.05, the numbers of up- and down-regulated differentially expressed genes (DEGs) were 617 and 561 at 0.5 days, 1475 and 1420 at 1 day, 827 and 736 at 2 days, and 1218 and 1307 at 5 days (Figure 6A and 6B). Importantly, even with a more stringent criteria of |log_2_ (fold change) |>1 and *p*_adj_ < 0.05, a substantial number of DEGs were identified at all time points, and the number of up/down-regulated DEGs were 539/117, 1098/1316, 599/577, 872/897, respectively (Figure S2). These findings indicate that GvmR has a broad impact on large sets of genes. Figure 6B provides detailed information on the number of genes showing consistent expression changes across various time points. We found that only two genes were significantly upregulated, while nine were significantly downregulated at all four time points. Among the downregulated genes, *scn3360* encodes a Xre family transcriptional regulator. A PSI-BLAST analysis to identify SCN3360 orthologs (with query coverage 90% to 100% and percent identity 60% to 100%) resulted in a total of 1395 hits, with over 99% of them originating from *Streptomyces* (as of sep. 2023), suggesting that SCN3360 orthologs are highly conserved in *Streptomyces*, and GvmR significantly affects the expression of some conserved regulatory genes. KEGG pathway enrichment analysis (|log_2_ (fold change) | > 0.5 and *p*_adj_ < 0.05) showed that the DEGs are mostly enriched in pathways, including the biosynthesis of secondary metabolites, ribosome, oxidative phosphorylation, biosynthesis of cofactors, and purine metabolism (Figure S3). We then focused on analysis of the transcriptional profiles of genes participating in ribosome, oxidative phosphorylation, and purine biosynthesis. Ribosomes serve as cellular factories for protein synthesis, with various ribosomal proteins comprising the principal components of the ribosome. In ΔgvmR, a large number of genes encoding ribosomal proteins exhibited significant downregulation at 0.5 days, followed by upregulation thereafter (Figure 6C, Table S3). Our unpublished data indicated that the cell biomass of ΔgvmR was lower than that of NEAU6 during the early stages of fermentation but became comparable after fermentation completion. This observation suggests that the inactivation of *gvmR* has the potential to impede cell growth during fermentation. Furthermore, AMP and ATP serve as precursors and energy sources, respectively, for guvermectin biosynthesis. At the middle fermentation stage (5 days), when NEAU6 produced guvermectin in abundance and ΔgvmR exhibited almost negligible guvermectin, most of the genes involved in oxidative phosphorylation process and purine synthesis showed a significant downward trend in ΔgvmR (Figure 6C and S4; Table S4 and S5), suggesting that GvmR may control guvermectin production by influencing the expression of genes associated with precursor and energy supply pathways.

**Figure 6.**
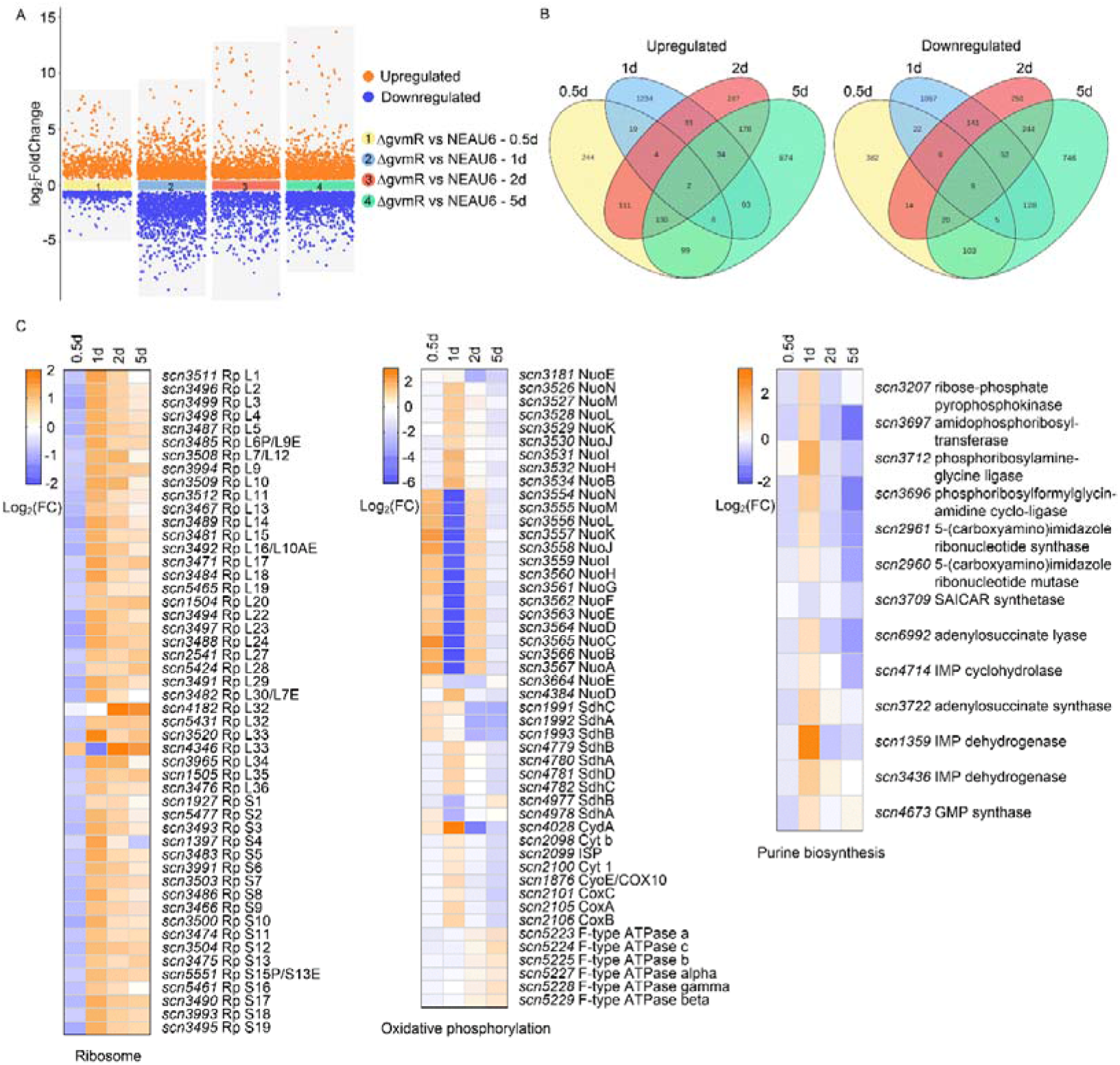
Transcriptome analysis of strains NEAU6 and ΔgvmR. A, scatter plots showing the overall distribution of DEGs identified via comparative transcriptome analysis between NEAU6 and ΔgvmR at different time points. B, the venn diagrams showing the numbers of DEGs identified in samples of NEAU6 and ΔgvmR. C, heat maps of genes involved in oxidative phosphorylation, ribosome, and purine biosynthesis. DEGs mentioned here are the ones that show significant expression change with the criteria |log_2_ (fold change) | > 0.5 and *p*adj < 0.05.

### Mining novel transcriptional regulators affecting guvermectin production from GvmR disturbed regulatory network

The *S. caniferus* NEAU6 genome encompasses approximately 652 transcriptional regulatory genes constituting 7.8% of the total chromosome genes (652/8435), excluding genes encoding RNA polymerase core enzymes and sigma factors. After inactivation of *gvmR*, the transcript levels of 252 transcriptional regulatory genes exhibited significant alterations at least at one time point (|log_2_ (fold change) | > 1 and *p*adj < 0.05) (Figure 7A), suggesting that GvmR has wide-ranging effects on expression of many transcriptional regulatory genes. Given the substantial changes in the expression of regulatory genes after the inactivation of *gvmR*, the question arises as to whether some of these genes also influence guvermectin production. To explore this possibility, we selected a total of 48 regulatory genes based on the significance criteria |log_2_ (fold change) |>2 for overexpression to assess their influence on guvermectin production (Figure 7A and 7B; Table S6). *scn5806*, a TetR family regulatory gene exhibiting high similarity to the autoregulator receptor (*arpA*) component of the quorum-sensing system, showed a significance degree of 1.0 < |log_2_ (fold change) | < 2.0 at 0.5 and 1 days (Table S6), was also chosen for overexpression. In the design of overexpression experiments, regulatory genes with the highest FPKM value ≥500 at least at one of the four time points were driven by their native promoters, while those with the highest FPKM value < 500 were controlled by the constitutive P_hrdB_. These gene overexpression plasmids were constructed using the integrative pSET152 and were further integrated into the ФC31 *attB* site of NEAU6. The resulting overexpression strains, along with the control strains NEAU6 and NEAU6/pSET152 were fermented and tested for guvermectin production. Despite the integration of pSET152 into NEAU6 leading to a decrease in guvermectin production, the experimentation yielded noteworthy results. Six out of the 49 regulatory genes exhibited significant effects on guvermectin production. Five of these genes (*scn3360*, *scn4836*, *scn4952*, *scn4970* and *scn6388*) had a positive effect while one (*scn5806*) demonstrated a negative influence (Figure 7C). Of the six regulatory genes, *scn3360* and *scn4952* encode Xre family regulators. SCN3360 orthologs are widespread in *Streptomyces*, while SCN4952 orthologs are only found in 23 *Streptomyces* genomes (Table S7). Xre family transcriptional regulators are ubiquitous across the three domains of life: eubacteria, archaea, and eukarya. Although *Streptomyces* genomes contain numerous Xre regulatory genes, only a few of these regulators, such as BldD, WhiJ, SCO4441 and CanR1, have been characterized to be involved in developmental differentiation and secondary metabolism (Romero-Rodríguez et al., 2015; Tian et al., 2020). It is worth noting that the ortholog of SCN3360, SCO4441 from *S. coelicolor*, shares 69% identity with SCN3360 and has been proved to activate diverse NPs production in various *Streptomyces* species (Santamaría et al., 2018). Here, the overexpression of *scn3360* significantly improved guvermectin production, further confirming *scn3360* and its orthologs are promising regulatory tools for promoting NP production.

**Figure 7.**
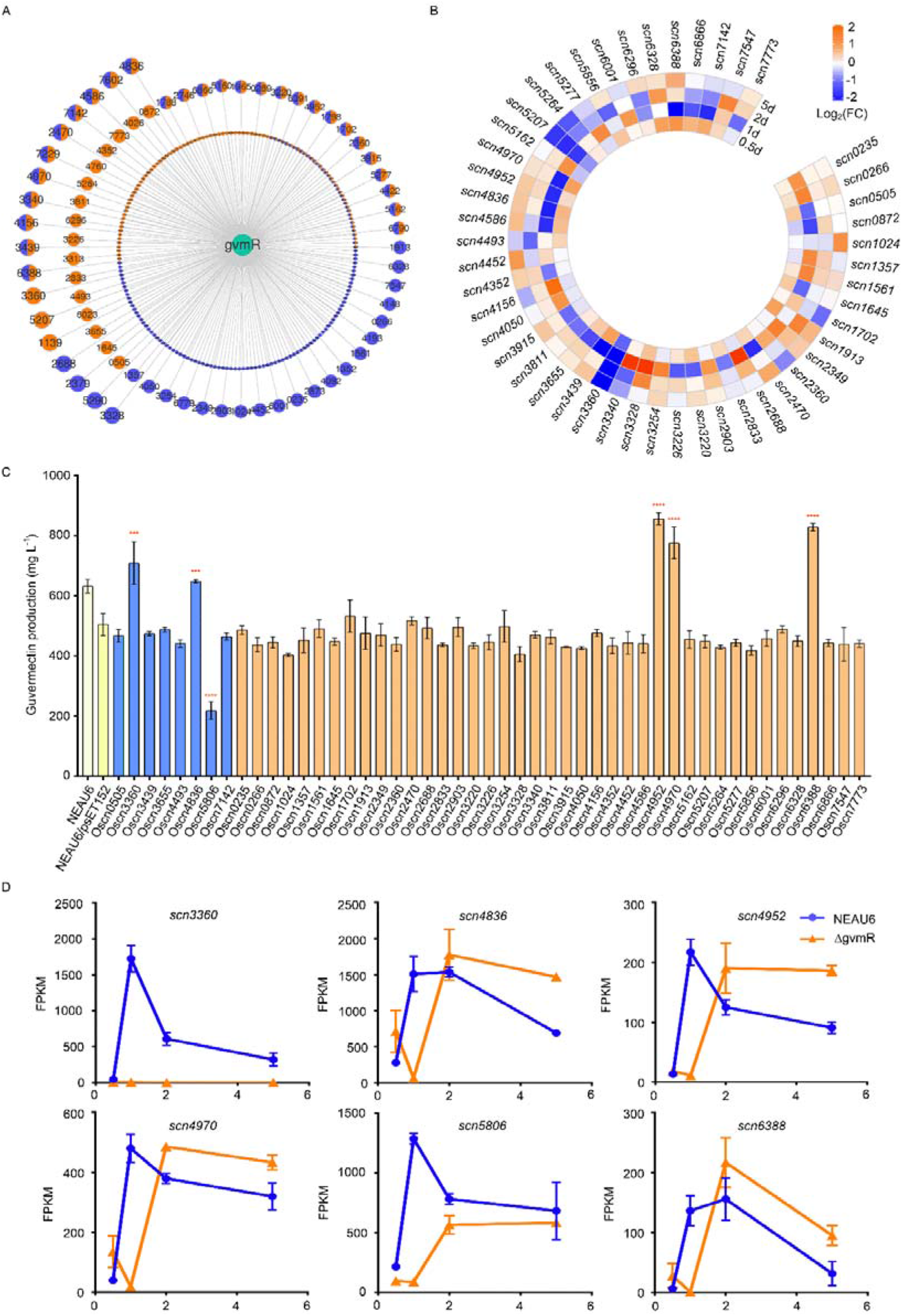
Influence of GvmR toward various transcriptional regulatory genes and identification of new regulators affecting guvermectin production. A, effects of GvmR on the expression of 252 transcriptional regulatory genes. Orange circles: regulatory genes that are significantly upregulated at 3 and/or 6 days; Blue circles: regulatory genes that are significantly downregulated at 3 and/or 6 days; bicolor circles: regulatory genes that are significantly upregulated at one detected time point and significantly downregulated at the other time point. Innermost circle: genes showing significant expression change with the criteria 1.0 < |log_2_ (fold change)| < 2.0; middle circles: genes showing significant expression change with the criteria 2.0 < |log_2_ (fold change) | < 4; Outermost circle: genes showing significant expression change with the criteria |log_2_ (fold change) | > 4. B, heatmap analysis of regulatory genes with the expression change criteria |log_2_ (fold change) | > 4 in ΔgvmR, and with FPKM > 50 at least at one time point in NEAU6 or ΔgvmR. C, gene overexpression to screen regulatory genes affecting guvermectin production. \*\*\**P* < 0.001, \*\*\*\**P* < 0.0001. D, effects of GvmR on the expression of new guvermectin regulatory genes.

*scn4836* encodes a DeoR family regulator whose orthologs are widespread in *Streptomyces*. Currently, only one ortholog of SCN4836, SdrA from *S. avermitilis*, has been reported as an activator of avermectin production and a repressor of oligomycin and filipin production. However, the corresponding regulatory mechanisms remain unclear. As an ArpA homolog, *scn5806* exhibits high similarity to the well-characterized autoregulator receptor AvaR1 of *S. avermitilis* (46% identity), ArpA (44% identity) of *S. griseus*, and ScbR of *S. coelicolor* (39% identity). ArpA and its homologs have been reported to modulate the production of NPs in various *Streptomyces* species by directly or indirectly controlling the expression of BGCs, and genetic manipulation of ArpA homologs has been demonstrated as a viable strategy for NP activation and overproduction (Tan et al., 2015). Notably, in this study, *scn5806* overexpression inhibited guvermectin production, suggesting that deletion of *scn5806* could potentially enhance guvermectin production. Additionally, *scn4970* encodes a regulator belonging to the GntR family, while scn6388 encodes a regulator containing a DUF955 domain. Despite their high conservation in *Streptomyces*, their functions have not been previously reported. Our findings indicate that overexpression of the two regulators can promote guvermectin production, suggesting their potential utility as practical tools for the efficient production of NPs. These results clearly prove that novel transcriptional regulators affecting guvermectin production can be mined from GvmR disturbed transcriptional regulatory network, providing more selectivity for construction of guvermectin high-producing strains.

### The complex differential control of GvmR toward the newly discovered guvermectin regulators

To determine the effects of GvmR on these newly identified guvermectin regulatory genes, we extracted their FPKM values from the transcriptomic data of NEAU6 and ΔgvmR for further comparative analysis. As shown in Figure 7D, after *gvmR* inactivation, the transcription profile changes of the six transcription regulators were categorized into two distinct groups. For *scn3360* and *scn5806*, transcription levels in ΔgvmR were markedly lower than those in NEAU6 at 1 and 2 days, and slightly lower than or equivalent to those of NEAU6 at 0.5 and 5 days, indicating a positive role of GvmR toward them. In contrast, the other four regulators exhibited lower transcription levels at 1 day but higher levels at 5 days in ΔgvmR compared to NEAU6. During this sampling period, the inactivation of *gvmR* appeared to delay the expression changes of these four genes, indicating a growth-stage-dependent differential effect of GvmR on them. This regulatory characteristic of delayed gene expression has been previously observed in *S. coelicolor*, in which, overexpressing *actII-ORF4*, a CSR gene of the actinorhodin BGC, first decreased and then increased transcription of the calcium-dependent antibiotic BGC relative to the control strain at 28 and 48 h respectively (Huang et al., 2005). EMSAs were then used to determine the regulatory relationships between GvmR and the promoter regions of these novel guvermectin regulatory genes. However, no binding was detected, suggesting that GvmR may indirectly affect the expression of these newly identified regulatory genes.

## DISCUSSION

The Biosynthesis of NPs in *Streptomyces* is intricately controlled by a sophisticated cellular regulatory network. with transcriptional regulators serving as pivotal framework components. These regulators play a crucial role in precisely modulating the transcription levels of target genes, thereby controlling the initiation and final titer of NPs (Liu et al., 2013). Transcriptional regulators with global or pleiotropic effects have the potential to coordinate the production of target NPs by simultaneously controlling the expression of multiple biological processes genes(Deng et al., 2022; Huang et al., 2005). Therefore, exploring titer-related factors from altered changed gene expression profiles influenced by transcriptional regulators is crucial for systematically understanding the regulatory network of NP biosynthesis and constructing high-titer strains. This study focuses on *gvmR*, an essential positive regulatory gene within the guvermectin cluster. Employing a multidisciplinary approach that includes genetics, biochemistry, and omics analysis, we demonstrate that GvmR is a pleiotropic regulator. It orchestrates guvermectin production by synergistically controlling BGCs, as well as genes associated with precursors and energy supply pathways, along with transcriptional regulatory networks (Figure 8).

**Figure 8.**
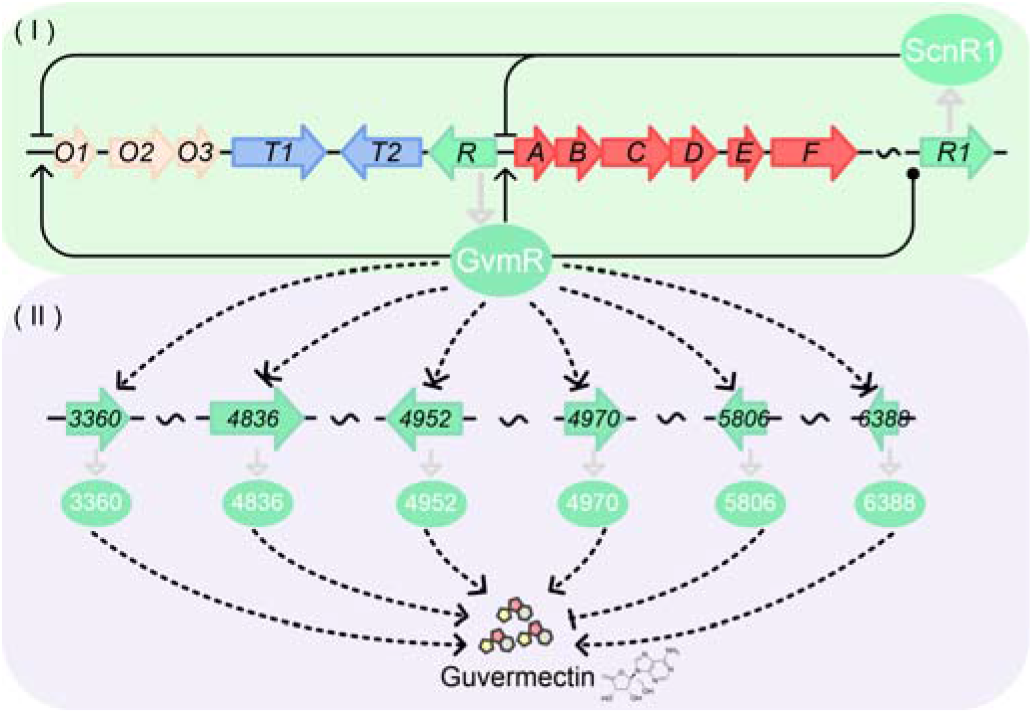
A plausible model for GvmR-centered regulation of guvermectin production in *S. caniferus* NEAU6. (I) GvmR is essential for guvermectin production. It directly activates transcription of the guvermectin cluster genes by binding to the promoter regions of *gvmR*, *gvmA* and *O1*. GvmR also directly binds to the promoter region of *scnR1*, another high conserved LacI-like regulatory gene, whose overexpression represses guvermectin production by binding to the promoter regions of *gvmR*, *gvmA* and *O1*. (II) GvmR indirectly activates transcription of *scn3360* and *scn5806* and dually influences transcription of *scn4836*, *scn4952*, *scn4970* and *scn6388*. Interestingly, five of them positively controls guvermectin production respectively, while *scn5806* represses guvermectin production. The cross-regulation between GvmR and ScnR1, and the differential disturbance of GvmR toward the other six regulators shed new light on the understanding of CSR’s regulatory pattern in coordinating NP production.

CSRs are bottom switches directly turning on or off expression of BGCs. Traditionally, CSR genes were thought to exclusively impact a limited set of genes, primarily the cognate BGC genes. In contrast, global/pleiotropic regulators located outside the BGCs were considered upper-level regulators that modulate CSR expression through regulatory cascades, thereby achieving control over multiple disparate NP BGCs (Huang et al., 2005). As the identification and characterization of CSRs increase and omics methods become more prevalent in the functional investigation of transcription regulators (Huang et al., 2005; Lin et al., 2020), it is evident that some CSRs exhibit a pleiotropic regulatory role, exerting extensive influence on the expression of genes associated with various BGCs, primary metabolism, and even morphological differentiation. For example, the overexpression of the *red* cluster activator gene, *redZ* from *S. coelicolor* could transiently increase transcription of *cda*, *red* and *act* clusters(Huang et al., 2005). Inactivation of *kelR*, a positive regulatory gene within a type II PKS cluster in *S. bingchenggensis*, not only leads to the absence of cognate compound but also results in nearly negligible expression of several other BGCs, including the milbemycin BGC(Wang et al., 2022). The key CSR of the nemadectin cluster, *nemR*, *in S. cyaneogriseus*, also directly controls four non-nemadectin BGC genes involved in precursor supply, morphological development and chemical secretion(Li et al., 2019). Consistent previous reports, we have also discovered that GvmR is a pleiotropic regulator affecting the expression of the guvermectin BGC and metabolic pathway genes necessary for guvermectin production. Interestingly, as a CSR, GvmR also establishes close interactive relationships with numerous transcriptional regulator genes. GvmR binds to the promoter region of *scnR1*, another LacI family regulatory genes conserved in *Streptomyces*. ScnR1, in turn, inhibits guvermectin production by binding to the same promoters as GvmR. Additionally, GvmR significantly disturbed the expression of 252 transcriptional regulatory genes, as revealed by comparative transcriptome analysis. At least four of the 48 transcription regulators with a substantial expression change (|log_2_ (fold change)| > 2) were verified to affect the production of guvermectin. Furthermore, five of the six newly discovered guvermectin regulators are widely existed in *Streptomyces*, underscoring the role of GvmR in modulating guvermectin biosynthesis by influencing the expression of several highly conserved regulators. This expands our comprehension of the regulatory patterns governed by CSR in the control of natural product (NP) biosynthesis. Of particular note, ScnR1, SCN4970 and SCN6388 are not only highly conserved, but also lack verified functional annotations. Many highly conserved transcriptional regulators were reported to exert differential control toward different NP biosynthesis (Li et al., 2023; Yang et al., 2022). Taking ScnR1 as an illustrative example, despite its role in repressing guvermectin production, our unpublished data reveal that its overexpression in *S. coelicolor* significantly enhances ACT biosynthesis. Hence, these newly discovered NP transcriptional regulators were candidate valuable genetic elements for constructing optimal microbial cell factories using synthetic biology approaches.

The identification of transcriptional regulators influencing NP production is a fundamental prerequisite for deciphering the biosynthetic regulatory network and achieving high-titer productions. CSRs are major regulators controlling expression abundance of NP BGCs, so effective reinforcement of BGCs can be achieved by manipulation of CSRs(Zhou et al., 2020). However, the generation of an optimal high-titer strain necessitates concurrent optimization of various physiological processes, encompassing precursor availability, energy supply, nutrient uptake and utilization, product efflux, and mycelium morphology (Lu et al., 2016; Ye et al., 2022). Therefore, it is imperative to explore novel transcription regulators that impact the final output from multiple perspectives, a crucial step toward comprehensively enhancing the production performance of NP producers. Luckily, advances of bioinformatics and high-throughput sequencing technologies facilitate the rapid acquisition of bacterial genome sequences, NP BGC information, and culture-dependent genome-wide gene expression profiles, thus providing new opportunities for devising strategies to unearth novel transcriptional regulators. Leveraging sequence information from target BGCs and the native producer’s genome enables the swift screening of candidate regulators through literature review and protein blast analysis. For instance, various *Streptomyces* strains utilize quorum-sensing systems like ArpA/AfsA or ArpA/Aco to regulate secondary metabolism (Zhang and Tan, 2023). Manipulation of such systems in new NP producers is likely to influence the final titer of the target NP. Regarding the utilization of transcriptome data, the Wen group conducted a comprehensive analysis by comparing the transcriptomes of the avermectin wild-type strain ATCC31267 with the overproducing strain 76-02-e. They selected transcriptional regulators exhibiting the highest expression changes for subsequent genetic validation, ultimately pinpointing multiple novel regulators influencing avermectin production (Guo et al., 2013). However, it is noteworthy that the generation of the overproducing strain 76-02-e involved a conventional random mutagenesis approach, which is not only time-consuming but also labor-intensive. Therefore, the development of a comparative omics-dependent strategy for mining transcriptional regulators, without relying on random mutagenesis to produce high-titer strains, is imperative. CSRs are easy to identify as they situate in NP BGCs. Some CSRs cross-regulate large set of other metabolic pathways genes situated far from BGCs. This suggests their potential role in coordinating the production of corresponding natural products through various physiological processes (Li et al., 2019). However, the regulatory roles of CSR toward various transcriptional regulatory genes attracted little attention, let alone the potential control of CSR on titers through its interaction with transcriptional regulators. Here, 49 transcriptional regulatory genes exhibiting significant expression changes following *gvmR* inactivation were carefully selected for overexpression experiments. Remarkably, five of these genes demonstrated positive regulatory effects on guvermectin biosynthesis. This suggests that it is indeed feasible to screen for new natural product regulators from the CSR perturbed transcriptional regulator network. CSR-engineered strains are easier to generate than those obtained through random mutagenesis. Consequently, this comparative transcriptome analysis between CSR-modified and the parental strains introduces a novel approach for the discovery of previously unrecognized NP transcriptional regulators. This methodological advancement holds the potential to accelerate the construction of high-titer strains based on regulatory principles.

The inactivation of *gvmR* had a profound impact on the expression of a substantial number of transcriptional regulatory genes, with a noteworthy effect observed in at least 252 genes. For genetic analysis, we focused our attention on a subset of 48 genes. Notably, an additional 204 genes exhibited moderate changes in expression (1 < |log_2_ (fold change) | < 2) but have not undergone comprehensive functional scrutiny. This implies the existence of potentially undiscovered transcriptional regulators that could play a role in influencing guvermectin production, warranting further investigation for validation. Due to the complex intertwined regulatory network and the versatility of transcriptional regulators, simple combinational manipulation of beneficial regulatory genes often fails to achieve the expected titer (Li et al., 2021). When more and more guvermectin biosynthesis regulators are discovered, how to combine and fine-tune these targets to construct a multifunctional high-titer module becomes a new challenge. Transcriptional control is the first step in modulating gene expression, particularly in bacteria. RNA-seq can be used to detect the abundance and dynamic changes of all genes in the genome under specific culture conditions. Performance of RNA-seq experiments toward various high-titer strains generated via engineering of regulatory genes will obtain multiple genomic gene expression profiles; then a comprehensive comparison of similarities and differences among sets of transcriptome profiles may provide a reference for further combination of beneficial targets.

## MATERIALS AND METHODS

### Strains, plasmids, primers, and culture conditions

The strains and plasmids used in this work were listed in Table S8; primers were summarized in Table S1. *Streptomyces caniferus* NEAU6 is a wild type guvermectin producer; ΔgvmR is a *gvmR* inactivation mutant generated from *S*. *caniferus* NEAU6 (Liu et al., 2023). *Streptomyces coelicolor* 1146 was used for GUS assays. *Escherichia coli* JM109 was used as a general host for propagating plasmids. *E. coli* ET12567 (pUZ8002) was used for transferring DNA from *E. coli* to *Streptomyces* by conjugation (Kieser et al., 2000). *E. coli* BL21 (DE3) was used to express GST-tagged GvmR and ScnR1. *S*. *caniferus* NEAU6 and its derivatives were cultured at 28°C on YMS agar medium for spore collection (Ikeda et al., 1988). Seed medium (2% sucrose, 2% maltdextrin, 6% soybean powder, and 0.3% CaCO_3_, pH 7.0) and fermentation medium (1% sucrose, 6% maltdextrin, 4% soybean powder, and 0.4% CaCO_3_, pH 7.2) were used for NEAU6 vegetative mycelium preparation and guvermectin production, respectively. AS-1 medium was prepared for GUS assays as described previously(Zhang et al., 2016). LB medium was used to cultivate *E. coli* strains. MS medium was used for conjugation from *E. coli* to *Streptomyces*(Kieser et al., 2000).

### Gene complementation and overexpression

For the complementation of *gvmR* in ΔgvmR, two plasmids pIJ10500::gvmR and pIJ10500::P_hrdB_gvmR were constructed. To construct pIJ10500::gvmR, a 1681-bp fragment containing the ORF and the upstream region of *gvmR* was amplified by PCR from *S. caniferus* NEAU6 genomic DNA using primers CNgvmR-F/R. This fragment was then inserted into the SpeI/XhoI sites of pIJ10500 by Gibson assembly to generate pIJ10500::gvmR. To construct pIJ10500::P_hrdB_gvmR, The *hrdB* promoter and the *gvmR* ORF were individually amplified from the genomic DNAs of *S. coelicolor* and *S. caniferus* NEAU6 with primer pairs PhrdB-F/R and CHgvmR-F/R. The obtained two fragments and the SpeI/XhoI digested pIJ10500 were ligated together through Gibson assembly to generate pIJ10500::P_hrdB_gvmR. The two complementation plasmids verified by PCR and sequencing were individually introduced into ΔgvmR by conjugal transfer to generate the complementation strains ΔgvmR/gvmR and ΔgvmR/P_hrdB_gvmR.

For the overexpression of *scnR1* in NEAU6, a 1131-bp fragment containing *scnR1* ORF was amplified from NEAU6 genomic DNA using primers H1544-F/R. This fragment was then cloned into the XbaI/EcoRI sites of pSET152::P_hrdB_, generating *scnR1* overexpression plasmid pSET152::P_hrdB_scnR1, which was further introduced into NEAU6 to obtain overexpression strain OscnR1.

To screen targets influencing guvermectin production from the 252 transcriptional regulatory genes that showed significant differential expression changes after *gvmR* inactivation, a variety of regulatory gene overexpression strains were constructed. For construction of *scn5806* overexpression strain, a 1205-bp fragment containing the ORF and the upstream region of *scn5806* was amplified from NEAU6 genomic DNA using primers N5806-F/R. This fragment was then cloned into the XbaI/EcoRI sites of pSET152, generating overexpression plasmid pSET152::scn5806. The construction processes of other regulatory genes was similar to that of constructing pSET152::scn5806 or pSET152::P_hrdB_scnR1, respectively. These overexpression plasmids were individually verified by PCR and sequencing and further introduced into NEAU6 by conjugal transfer to generate overexpression strains as listed in Table S8.

### HPLC analysis of guvermectin

For extraction of guvermectin, 0.4 ml of whole cell fermentation broth was mixed with four-fold volume of ethanol and treated by ultrasound for 1 h, followed by centrifugation at 12000 rpm for 10 min. The resulting supernatant was used for HPLC analysis. HPLC analysis was carried out by Agilent 1260 II system with an Sq-C18 column (Zorbax, 4.6×250 mm, 5 μm) at a flow rate of 0.8 ml/min with 90% solvent A and 10% solvent B in 30 min (Solvent A: ddH_2_O; Solvent B: CH_3_CN) and detected at 260 nm.

### RNA preparation, RT-PCR, and qRT-PCR

Total RNAs were extracted from NEAU6 and its derivatives fermented for 0.5, 1, 2 and 5 days. RNA isolation, examination of RNA quality and quantity, cDNA synthesis, RT-PCR and qRT-PCR were performed as previous report (Zhang et al., 2013).

### Expression and purification of GvmR and ScnR1

To construct GvmR expression plasmid, the coding region of *gvmR* was amplified from the genomic DNA of NEAU6 using primer pair GEXgvmR-F/R. The PCR product was cloned into BamHI/XhoI digested pGEX-4T-1 to obtain pGEX-4T-1::gvmR. The construction process of ScnR1 expression plasmid (pGEX-4T-1::scnR1) was similar to that of constructing pGEX-4T-1::gvmR. The overexpression and purification processes were the same as described previously(Li et al., 2019).

### Electrophoretic mobility shift assays (EMSAs)

EMSAs were performed as reported previously(Zhang et al., 2013). The promoter probes were amplified from the genomic DNA of NEAU6 by PCR with primer pairs listed in Table S1.

### Site-directed mutagenesis of GvmR and ScnR1 binding sequences

To evaluate the specificity of GvmR on its binding sequences, the bidirectional promoter region P_R-A_ was inserted into EcoRV-digested pBluescript KS (+) to generate pBlu::P_R-A_. Three fragments containing mutated P_R-A_ were amplified using pBlu::P_R-A_ as template by primer pairs NheI-F/R, NdeI-F/R and NheI-F/NdeI-R, respectively, which were self-ligated, generating mutant plasmids pBlu::P_R-A_-M1, pBlu::P_R-A_-M2 and pBlu::P_R-A_-M3. These three plasmids were further used as templates separately to amplify mutant probes P_R-A_-M1, P_R-A_-M2 and P_R-A_-M1 by primer pairs gvmRA-pF/R. The binding activities of GvmR or ScnR1 to the three probes was subsequently determined by EMSAs.

### GUS assays

GUS activities were measured as described previously(Myronovskyi et al., 2011; Sherwood and Bibb, 2013). To determine the potential promoter activities of *gvmT1*, *gvmT2*, *gvmE* and *gvmF*, the corresponding promoters of them were obtained from the NEAU6 genomic DNA by PCR with primer pairs gvmT1GUSA-pF/R, gvmT2GUSA-pF/R, gvmEGUSA-pF/R, gvmFGUSA-pF/R, respectively (Table S1), which were cloned into the upstream of *gusA* reporter based on pSET152 by Gibson assembly, generating plasmids pSET152::P_T1_gusA, pSET152::P_T2_gusA, pSET152::P_E_gusA, pSET152::P_F_gusA. These four plasmids together with the control plasmid pSET152::gusA were introduced into M1146 by conjugal transfer to generate derivatives M1146/gusA, M1146/P_T1_gusA, M1146/P_T2_gusA, M1146/P_E_gusA and M1146/P_F_gusA, respectively.

To determine the importance of the 14-bp palindromic sequences for GvmR regulatory activity *in vivo*, the promoter region of *gvmA* (P_A_) and the corresponding mutant probes (P_A_-M1, P_A_-M2 and P_A_-M1) were PCR amplified using pBlu::P_R-A_, pBlu::P_R-A_-M1, pBlu::P_R-A_-M2 and pBlu::P_R-A_-M3 as templates by primers PgvmA*-* XF /R, respectively. Each of these four fragments, the promoter less *gusA* and P_hrdB_ activated *gvmR* were assembled into the PvuI/EcoRV digested pSET152::hyg, generating plasmids pSyg::P_A_gusA::P_hrdB_gvmR, pSyg::P_A_*-*M1gusA::P_hrdB_gvmR, pSyg::P_A_*-*M2gusA::P_hrdB_gvmR, pSyg::P_A_*-*M1gusA::P_hrdB_gvmR and pSyg::P_F_gusA::P_hrdB_gvmR. In addition, P_A_ and the promoter less *gusA* were inserted into the XbaI/EcoRV digested pSET152::hyg, generating control plasmid pSyg::P_A_gusA. These plasmids were then introduced into ΔgvmR by conjugation.

### RNA-seq analysis

RNA-seq was performed by Novogene Bioinformatics Technology Co. Ltd. Three independent biological replicates were analyzed at 0.5, 1, 2 and 5 days, Calculation of Gene expression levels and the threshold of *P*-value were the same as previous report (Ye et al., 2022). The DEGs were selected based on |log_2_ (fold change) | > 0.5 and *p*_adj_ < 0.05.

### Statistical analysis

All experiments were performed at three biological triplicates. Data were presented as averages of three triplicates. Significance was analyzed by Student’s *t*-test, and the significance were presented as follows, \*\*\*\**p* <0.0001, \*\*\**p* < 0.001.

### Bioinformatics analysis

The sequence alignment and domains of potential protein-coding sequences (CDSs) were documented by publicly available databases and their application tools, including Pfam (http://www.pfam.xfam.org/) and SMART (http://www.smart.embl-heidelberg.de/). Heat maps were created by the Graphpad software and OmicShare Tool (https://www.omicshare.com/tools/Home/Soft/pathwaygseasenior). VENN and scatter diagram analysis were created by OmicShare Tool (https://www.omicshare.com/tools/Home/Soft/pathwaygseasenior). Motif-based sequence analysis tools were creat by MEME (https://meme-suite.org/meme/index.html).

### Data availability

The genome sequence of *Streptomyces caniferus* NEAU6, and the raw datasets of comparative transcriptome between NEAU6 and ΔgvmR have been deposited in China National Microbiology Data Center (NMDC NMDC60139508 and NMDC NMDC40049731-49754).

## Compliance and ethics

*The author(s) declare that they have no conflict of interest*.

## Acknowledgements

This study was supported by grants from the National Natural Science Foundation of China (32030090 and 32272635). We are grateful to Prof. Mervyn Bibb (John Innes Centre, Norwich, UK) for providing S. coelicolor M1146 and Prof. Mark Buttner (John Innes Centre, Norwich, UK) for providing pIJ10500.

