## Supplementary material for "Omics-assisted systematic exploration of the intricate regulatory network of guvermectin biosynthesis centered by the cluster-situated regulator GvmR in *Streptomyces*": 20231120-Supplemental File

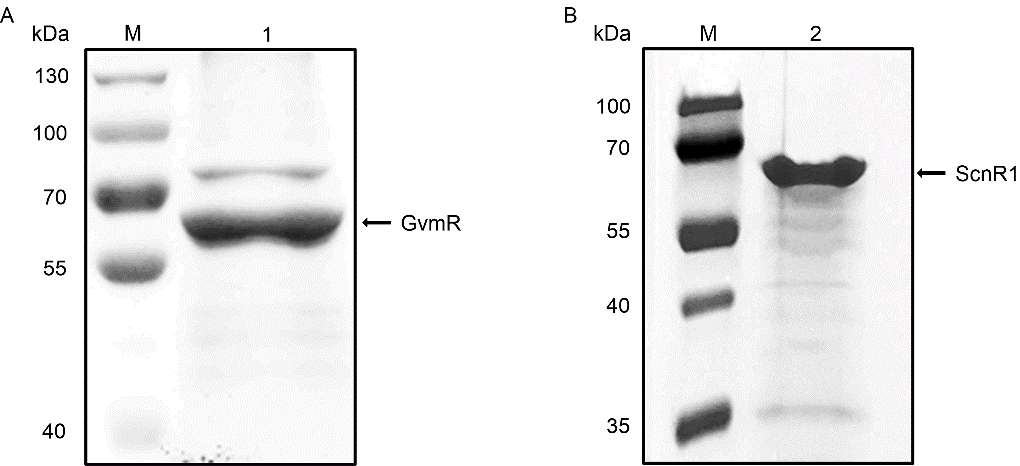

**Figure S1** SDS-PAGE analysis of soluble GST-tagged GvmR (A) and ScnR1 (B) proteins in *E. coli* BL21 (DE3). M, protein marker. 1, purified GST-tagged GvmR (63.5 kDa). 2, purified GST-tagged ScnR1 (64.8 kDa).

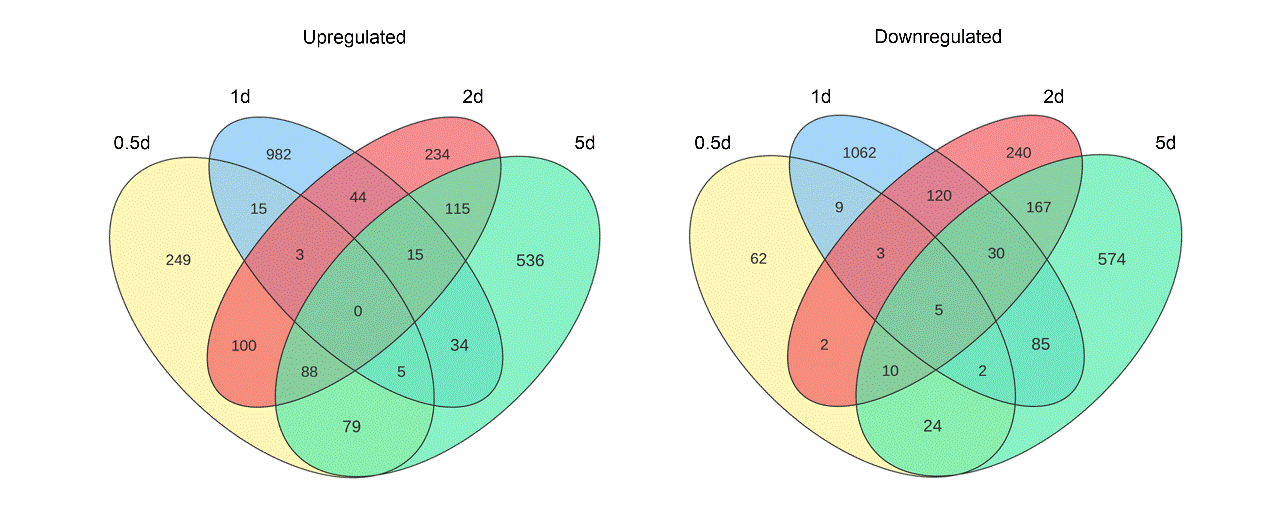

**Figure S2** The venn diagram showing the numbers of DEGs with the criteria of |log_2_(fold change)| > 1.0 and *p*adj < 0.05 in samples of NEAU6 and ΔgvmR.

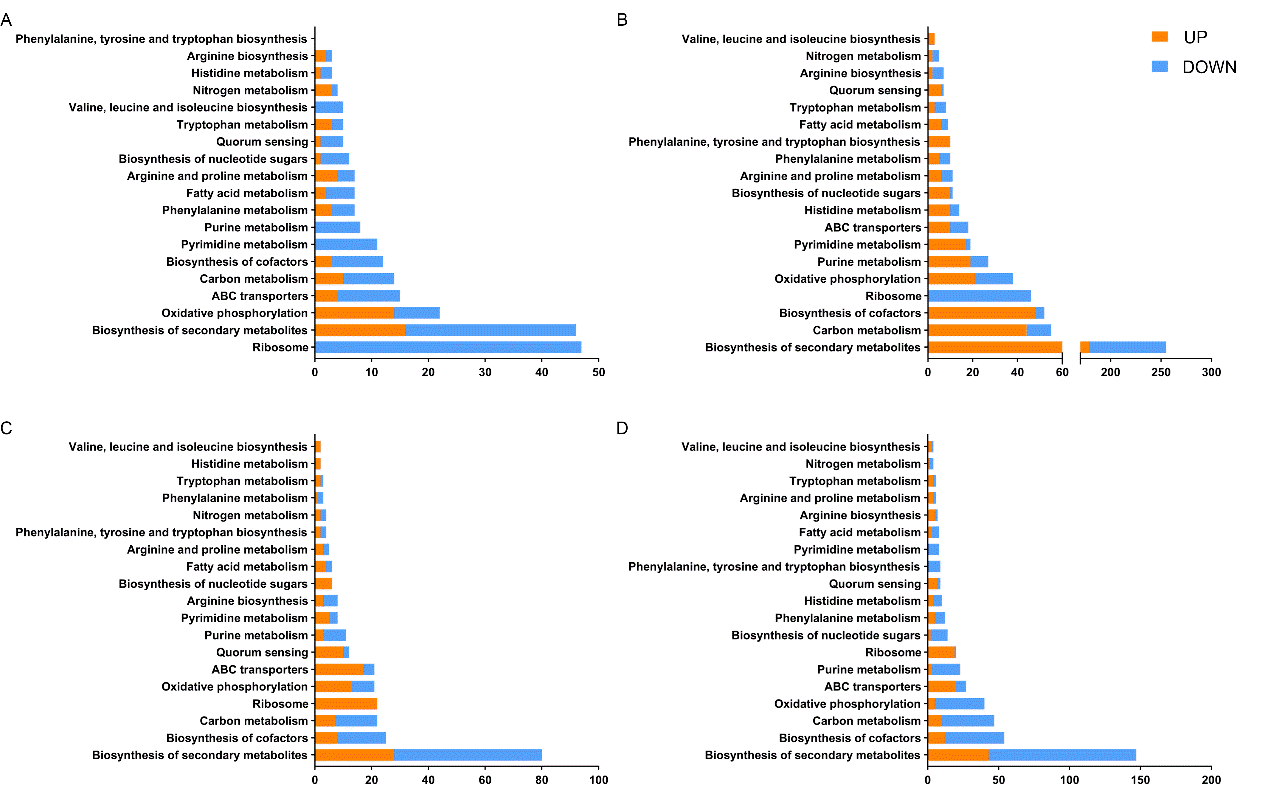

**Figure S3** Functional categories of annotated DEGs between strains *S. caniferus* NEAU6 and ΔgvmR based on the KEGG pathway analysis. A, 0.5 day. B, 1 day. C, 2 day. D, 5 day.

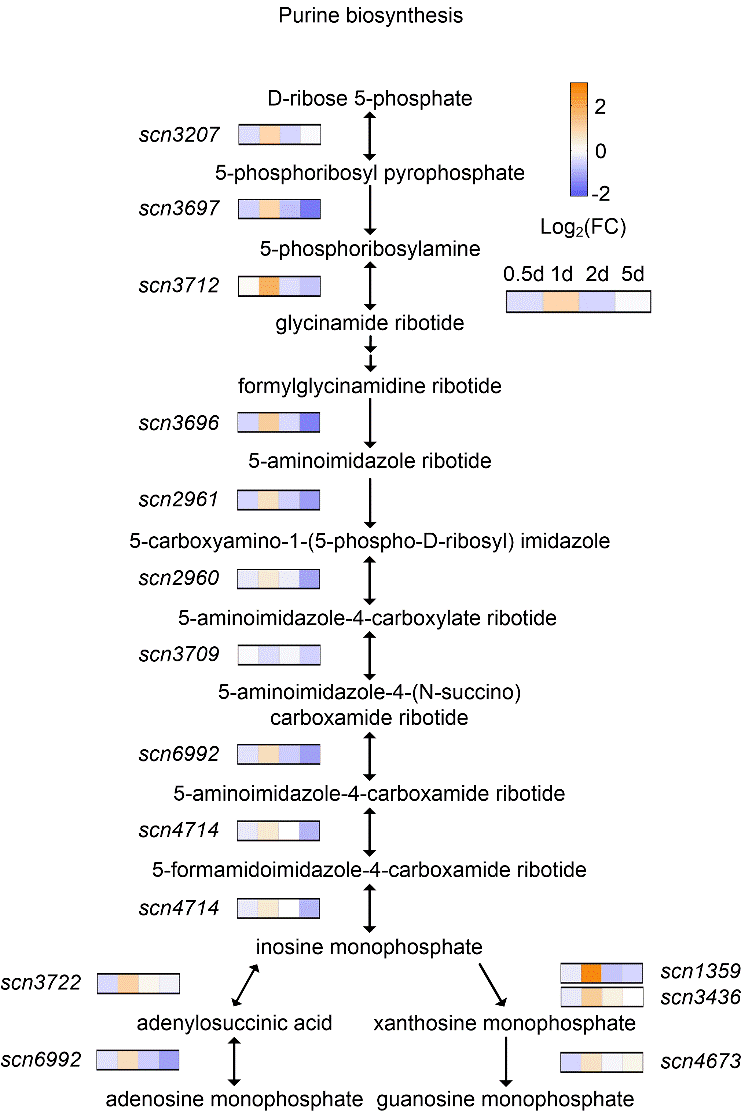

**Figure S4** The expression changes of genes involved in purine biosynthesis

**Table S1** Primers used in this work

| **Primers** | **Sequence (5'-3') ^a^** | **Usage** |
| --- | --- | --- |
| **For gene complementation and overexpression** | | |
| CNgvmR-F | **CTTGTAGTCGCCGTCGTGGTCCTTGTAGTC**CGCTGATGGCCAGGGCGAGGATT | Complementation of *gvmR* controlled by native promoter |
| CNgvmR-R | **TTTTTGGCCTTGAAATCGTTAGTTAGGCTA**CTCGAACAGTTCCTCCGCGGTGCC |  |
| CHgvmR-F | **AGTCGCCGTCGTGGTCCTTGTAGTC**TTAATTAAGGGCCTTCACCGCCGT | Complementation of *gvmR* controlled by *hrdB* promoter |
| CHgvmR-R | **GGCCTTGAAATCGTTAGTTAGGCTA**CCGCCTTCCGCCGGAACG |  |
| PhrdB-F | **AGTCGCCGTCGTGGTCCTTGTAGTC**TTAATTAAGGGCCTTCACCGCCGT | *hrdB* promoter |
| PhrdB-R | **GGCCTTGAAATCGTTAGTTAGGCTA**CCGCCTTCCGCCGGAACG |  |
| H1544-F | **TTTTCAACGTTCCGAGAGGTTGTTC**gtgaccagcaacgagaccaccac | Overexpression of *scnR1* |
| H1544-R | **TGCCAAGCTTGGGCTGCAGGTCGACTCTA**ggcgcttcgggcgaggac |  |
| H0235-F | **TTTTCAACGTTCCGAGAGGTTGTTC**ATGGCTTCCACACGGTCGTCGCT | Overexpression of *scn0235* |
| H0235-R | **TGCCAAGCTTGGGCTGCAGGTCGAC**GTCGCCGTCCCCTCTGCCG |  |
| H0266-F | **TTTTCAACGTTCCGAGAGGTTGTTC** GTGGAGCCTGACCACGTAGACG | Overexpression of *scn0266* |
| H0266-R | **TGCCAAGCTTGGGCTGCAGGTCGAC**AGGCCGATCTGCTCGTGCTG |  |
| N0505-F | **CAGGAAACAGCTATGACATGATTAC**ACCGAGCTGCCGAAAGGA | Overexpression of *scn0505* |
| N0505-R | **TGCCAAGCTTGGGCTGCAGGTCGAC**GGTGACCGGGGTGGAAGTAC |  |
| H0872-F | **TTTTCAACGTTCCGAGAGGTTGTTC**ATGTTTATGAACTCCTCGACGACCC | Overexpression of *scn0872* |
| H0872-R | **TGCCAAGCTTGGGCTGCAGGTCGAC**CCCTGGTGAGCGGTGAAACG |  |
| H1024-F | **TTTTCAACGTTCCGAGAGGTTGTTC**ATGCCCTCAGACGTGCCCCT | Overexpression of *scn1024* |
| H1024-R | **TGCCAAGCTTGGGCTGCAGGTCGAC**TGCTGGAAGTCGGGTCAGGGAT |  |
| H1357-F | **TTTTCAACGTTCCGAGAGGTTGTTC**GTGCGACTGAACGATCTCGACGA | Overexpression of *scn1357* |
| H1357-R | **TGCCAAGCTTGGGCTGCAGGTCGAC**CGGCCTCGTCACTGCGTATC |  |
| H1561-F | **TTTTCAACGTTCCGAGAGGTTGTTC**ATGGCACGCACCATCCAGTC | Overexpression of *scn1561* |
| H1561-R | **TGCCAAGCTTGGGCTGCAGGTCGAC**CCCCGTCAAGGGTTCCGT |  |
| H1645-F | **TTTTCAACGTTCCGAGAGGTTGTTC**GTGACCACCTTCGCTCCCG | Overexpression of *scn1645* |
| H1645-R | **TGCCAAGCTTGGGCTGCAGGTCGAC**GAAAAACGGCGGCAAAACCA |  |
| H1702-F | **TTTTCAACGTTCCGAGAGGTTGTTC**ATGTTGGAGACCTCGGCACGT | Overexpression of *scn1702* |
| H1702-R | **TGCCAAGCTTGGGCTGCAGGTCGAC**GTCCACGCTGCCCTTCCTCTA |  |
| H1913-F | **TTTTCAACGTTCCGAGAGGTTGTTC**ATGTCCGAGACCACGACGTATCT | Overexpression of *scn1913* |
| H1913-R | **TGCCAAGCTTGGGCTGCAGGTCGAC** CTCCGTGCATCTGAAGCTGTC |  |
| H2349-F | **TTTTCAACGTTCCGAGAGGTTGTTC**ATGGCGACCCCGACGCCGCATA | Overexpression of *scn2349* |
| H2349-R | **TGCCAAGCTTGGGCTGCAGGTCGAC**ACGGGAATGATCGCGTAGCGGTGGA |  |
| H2360-F | **TTTTCAACGTTCCGAGAGGTTGTTC**GTGTCTGAACCCGCATCGC | Overexpression of *scn2360* |
| H2360-R | **TGCCAAGCTTGGGCTGCAGGTCGAC**TCGAGCCAGGGAGTCAGGAA |  |
| H2470-F | **TTTTCAACGTTCCGAGAGGTTGTTC**ATGGCCAATGCCTCGCGACAA | Overexpression of *scn2470* |
| H2470-R | **TGCCAAGCTTGGGCTGCAGGTCGAC**GGAACTCGTGGGCAACCTCT |  |
| H2688-F | **TTTTCAACGTTCCGAGAGGTTGTTC**GTGAGTGTCACCTCGGACCCC | Overexpression of *scn2688* |
| H2688-R | **TGCCAAGCTTGGGCTGCAGGTCGAC**CCACGCATCTGGATTCCTGAC |  |
| H2833-F | **TTTTCAACGTTCCGAGAGGTTGTTC**ATGGCGATCGATCATTTGGACG | Overexpression of *scn2833* |
| H2833-R | **TGCCAAGCTTGGGCTGCAGGTCGAC**GACGTACTCCCAGAAGCTCACTCG |  |
| H2903-F | **TTTTCAACGTTCCGAGAGGTTGTTC**ATGGCAAAAGACGGCAGCGG | Overexpression of *scn2903* |
| H2903-R | **TGCCAAGCTTGGGCTGCAGGTCGAC**GCGGAGCACCACCGAAATG |  |
| H3220-F | **TTTTCAACGTTCCGAGAGGTTGTTC**GTGCAACGCATCCGGGTTCTGG | Overexpression of *scn3220* |
| H3220-R | **TGCCAAGCTTGGGCTGCAGGTCGAC**CCCACGACCGTCTGCCCCTC |  |
| H3226-F | **TTTTCAACGTTCCGAGAGGTTGTTC** ATGGGCGTGCGGCTCATGGT | Overexpression of *scn3226* |
| H3226-R | **TGCCAAGCTTGGGCTGCAGGTCGAC** AACTCCAGGCCACGACCGAGGAA |  |
| H3254-F | **TTTTCAACGTTCCGAGAGGTTGTTC**ATGGGACGCAGCCGGCTGACTC | Overexpression of *scn3254* |
| H3254-R | **TGCCAAGCTTGGGCTGCAGGTCGAC**TACGAGGGCTGCGGTGAAGGG |  |
| H3328-F | **TTTTCAACGTTCCGAGAGGTTGTTC**ATGGAACGGACACAGACGCT | Overexpression of *scn3328* |
| H3328-R | **TGCCAAGCTTGGGCTGCAGGTCGAC**TGAGGATCATGCGCGTACTACT |  |
| H3340-F | **TTTTCAACGTTCCGAGAGGTTGTTC**ATGATCTTCCTTTCCGGTTCCA | Overexpression of *scn3340* |
| H3340-R | **TGCCAAGCTTGGGCTGCAGGTCGAC**ACCCCCCTACTCCTGCCACT |  |
| N3360-F | **CAGGAAACAGCTATGACATGATTAC**AGGCGTTGGGCTGATTGC | Overexpression of *scn3360* |
| N3360-R | **TGCCAAGCTTGGGCTGCAGGTCGAC**GACGACGCTACGCTCCAACAT |  |
| N3439-F | **CAGGAAACAGCTATGACATGATTAC**ACCTCCTTCGCCGATGTCT | Overexpression of *scn3439* |
| N3439-R | **TGCCAAGCTTGGGCTGCAGGTCGAC**CCCCGAGGAACTCCATACC |  |
| N3655-F | **CAGGAAACAGCTATGACATGATTAC**CCGTCCTCACCCCGGCGTAT | Overexpression of *scn3655* |
| N3655-R | **TGCCAAGCTTGGGCTGCAGGTCGAC**CCACACTACCTACATCGGGCCA |  |
| N3811-F | **TTTTCAACGTTCCGAGAGGTTGTTC**GTGCTGGTGGCCGACGACCA | Overexpression of *scn3811* |
| N3811-R | **TGCCAAGCTTGGGCTGCAGGTCGAC**GCACACCAACGCCGTCTACTTCC |  |
| N3915-F | **TTTTCAACGTTCCGAGAGGTTGTTC**ATGACCATGGCCAAGCGCGA | Overexpression of *scn3915* |
| N3915-R | **TGCCAAGCTTGGGCTGCAGGTCGAC**ATCAGACCTGGTCAGCGACCGGA |  |
| H4050-F | **TTTTCAACGTTCCGAGAGGTTGTTC**ATGGCGCTTCAGGCGGCG | Overexpression of *scn4050* |
| H4050-R | **TGCCAAGCTTGGGCTGCAGGTCGAC**GGAGGAGAAAGGAACGGCGAAG |  |
| H4156-F | **TTTTCAACGTTCCGAGAGGTTGTTC**GTGGAGGAGCTCGTGGCCGC | Overexpression of *scn4156* |
| H4156-R | **TGCCAAGCTTGGGCTGCAGGTCGAC**TGTAGTTCCTTTTCATCCTGCGGGT |  |
| H4352-F | **TTTTCAACGTTCCGAGAGGTTGTTC**GTGCGGCGGATCCTGCTCGG | Overexpression of *scn4352* |
| H4352-R | **TGCCAAGCTTGGGCTGCAGGTCGAC**CGCCGTCGATGAACTGATACGCCAA |  |
| H4452-F | **TTTTCAACGTTCCGAGAGGTTGTTC**ATGACGCCCACGGACGCCGA | Overexpression of *scn4452* |
| H4452-R | **TGCCAAGCTTGGGCTGCAGGTCGAC**ATGACGCCCACGGACGCCGA |  |
| N4493-F | **CAGGAAACAGCTATGACATGATTAC**GGCGACAAACGGCGAGCAAC | Overexpression of *scn4493* |
| N4493-R | **TGCCAAGCTTGGGCTGCAGGTCGAC**CCAGTCTTGCCACATCATCCCG |  |
| N4586-F | **TTTTCAACGTTCCGAGAGGTTGTTC**ATGCTCGGGGCCGAGTTACG | Overexpression of *scn4586* |
| N4586-R | **TGCCAAGCTTGGGCTGCAGGTCGAC**ACCGTGCGGGTCCTTGGAGT |  |
| N4836-F | **CAGGAAACAGCTATGACATGATTAC**CGATGTACCACTTCCTCGGTGACGC | Overexpression of *scn4836* |
| N4836-R | **TGCCAAGCTTGGGCTGCAGGTCGAC**CGCCGCCGGGAGCCGTGTT |  |
| H4952-F | **TTTTCAACGTTCCGAGAGGTTGTTC**GTGGCCAAAGAAATTGATCCCT | Overexpression of *scn4952* |
| H4952-R | **TGCCAAGCTTGGGCTGCAGGTCGAC**GAGGCCGGTTATGAAGGTGG |  |
| H4970-F | **TTTTCAACGTTCCGAGAGGTTGTTC**GTGCCTTTCGGTGAGCAGCCC | Overexpression of *scn4970* |
| H4970-R | **TGCCAAGCTTGGGCTGCAGGTCGAC**TTGCCAACAAACCATCGTGAAAC |  |
| H5162-F | **TTTTCAACGTTCCGAGAGGTTGTTC**ATGCCCCCCGTCTTCGCCCA | Overexpression of *scn5162* |
| H5162-R | **TGCCAAGCTTGGGCTGCAGGTCGAC**GCGCTGCGGAAGTGCGTCGG |  |
| H5207-F | **TTTTCAACGTTCCGAGAGGTTGTTC**TTGTCCGGTCGGATCCTCGTT | Overexpression of *scn5207* |
| H5207-R | **TGCCAAGCTTGGGCTGCAGGTCGAC**GGGTCACAGGACAAGCGTATGG |  |
| H5264-F | **TTTTCAACGTTCCGAGAGGTTGTTC**ATGCCGGTGAACCTGCACTTC | Overexpression of *scn5264* |
| H5264-R | **TGCCAAGCTTGGGCTGCAGGTCGAC**CGAAGGTGGAGACCGTGGAGC |  |
| H5277-F | **TTTTCAACGTTCCGAGAGGTTGTTC**ATGGTCGCCCGCATGCAC | Overexpression of *scn5227* |
| H5277-R | **TGCCAAGCTTGGGCTGCAGGTCGAC**AGGTCTGAGCGGAGCGTGGC |  |
| N5806-F | **CAGGAAACAGCTATGACATGATTAC**CCGCAGGCGTGTTGTACCAGTT | Overexpression of *scn5806* |
| N5806-R | **TGCCAAGCTTGGGCTGCAGGTCGAC**CGGTGGGGGGAGGTGTCAGA |  |
| H5856-F | **TTTTCAACGTTCCGAGAGGTTGTTC**GTGTCTGTTCTCCTCGAGCAACCTT | Overexpression of *scn5856* |
| H5856-R | **TGCCAAGCTTGGGCTGCAGGTCGAC**TCGTGTCTGCTGCGGTCTTTT |  |
| H6001-F | **TTTTCAACGTTCCGAGAGGTTGTTC** ATGGGGCAGCACGACGCGCA | Overexpression of *scn6001* |
| H6001-R | **TGCCAAGCTTGGGCTGCAGGTCGAC**GTTGGAGGAGACCCGGAGCTGAGGC |  |
| H6296-F | **TTTTCAACGTTCCGAGAGGTTGTTC**ATGGATCTGCTGTCCCTGCGCTA | Overexpression of *scn6296* |
| H6296-R | **TGCCAAGCTTGGGCTGCAGGTCGAC**GCCGGCGCTCAAGAACACCT |  |
| H6328-F | **TTTTCAACGTTCCGAGAGGTTGTTC**GTGGAGGAGCGCATGGCGAA | Overexpression of *scn6328* |
| H6328-R | **TGCCAAGCTTGGGCTGCAGGTCGAC** GAACGTGTTCCATTTTCCACAAGCC |  |
| H6388-F | **TTTTCAACGTTCCGAGAGGTTGTTC**ATGCCTACCGCATTCCCCCAGG | Overexpression of *scn6388* |
| H6388-R | **TGCCAAGCTTGGGCTGCAGGTCGAC**AGTCCGAGCGCCGGGCCCTA |  |
| H6866-F | **TTTTCAACGTTCCGAGAGGTTGTTC** ATGGAGACCGGTGCCGATCC | Overexpression of *scn6866* |
| H6866-R | **TGCCAAGCTTGGGCTGCAGGTCGAC**CGGCCTCCCAATCGCTCT |  |
| N7142-F | **CAGGAAACAGCTATGACATGATTAC**CTCCGCACCCGCTTCCCTAT | Overexpression of *scn7142* |
| N7142-R | **TGCCAAGCTTGGGCTGCAGGTCGAC**GGCTGATGGGACCGTGTTGC |  |
| H7547-F | **TTTTCAACGTTCCGAGAGGTTGTTC**ATGACAAGAGTTCTGCTGATCGAGG | Overexpression of *scn7547* |
| H7547-R | **TGCCAAGCTTGGGCTGCAGGTCGAC**GAAGAGAAAGGCGACGACCAGA |  |
| H7773-F | **TTTTCAACGTTCCGAGAGGTTGTTC**GTGAGCACCTCGCTGCTCTAC | Overexpression of *scn7773* |
| H7773-R | **TGCCAAGCTTGGGCTGCAGGTCGAC**GCCACTACTGCCGCACTTCTACT |  |
| **For qRT-PCR** | | |
| real-16s-F | TGTCGTGAGATGTTGGGTTAAG | Control |
| real-16s-R | TCATTGTACCGGCCATTGTAG |  |
| real-T1-F | GGACTCCTCAGCATTCTTGT | Transcription analysis |
| real-T1-R | AGTGACCCACACCTGTTTC | Transcription analysis |
| real-T2-F | CCCATATGCCGATCCTGTC | Transcription analysis |
| real-T2-R | ATCACGGTGTTGATGGTCAG |  |
| real-R-F | TGGGTTATCGGCTCAATCTTTC | Transcription analysis |
| real-R-R | TGTGCACATCGGCGAAAT |  |
| real-A-F | ATGAACGCAAGGCACTGA | Transcription analysis |
| real-A-R | CATGTCAGGACCTCGAACAG |  |
| real-B-F | GGGACGACAACCATCCTG | Transcription analysis |
| real-B-R | GTAGTCGGACGCGAAGATG | Transcription analysis |
| real-C-F | TACCGCACCTCGCCTAT | Transcription analysis |
| real-C-R | GGTGATGAGCTGGTCGATAC | Transcription analysis |
| real-D-F | CGAACACGTCGCACAGTTA | Transcription analysis |
| real-D-R | AGGTCGCTGATGGTCTCC |  |
| real-E-F | TGATCGCCCGCAAGAAG | Transcription analysis |
| real-E-R | TCCTCACCGAGGAAGAGTT |  |
| real-F-F | TGACCGAGGTGGACCAT | Transcription analysis |
| real-F-R | GATGAAGATGTCGCCCAGTT |  |
| real-R1-F | GCCCTGACCACCGAATTCTT | Transcription analysis |
| real-R1-R | GGGTAGAGGACCACTCCGAA |  |
| **For protein expression** | | |
| GEXgvmR-F | GGAATTCTTGTCTGAAGGACTCCACAGGGCAGGGAACGCACC | Overexpression of GvmR in *E.coli* |
| GEXgvmR-R | CCGCTCGAGCCGCCGTGGCGGCCCCACGGAG |  |
| GEXscnR1-F | **GGTTCCGCGTGGATCCCCGGAATTC**GTGACCAGCAACGAGACCACCAC | Overexpression of ScnR1 in *E.coli* |
| GEXscnR1-R | **GTCAGTCACGATGCGGCCGCTCGAG**CTCCCCGCCGGGGCGCG |  |
| **For EMSAs** | | |
| gvmRA-pF | CCACTTTCCCGCCATCACCAA | Probe P_R-A_ |
| gvmRA-pR | CGTACCGCACGAGGCTCTGC |  |
| gvmT1-pF | GATGGTGTACGCGATGATGATGC | Probe P_T1_ |
| gvmT1-pR | ACGAAGGCGACCAGCAGGAA |  |
| gvmT2-pF | GGCGTAGCCGGAGACCAGGTA | Probe P_T2_ |
| gvmT2-pR | GGTCATCTGCGACGACGACAATCT |  |
| gvmE-pF | GTGATGAGCGTCGAACCGGG | Probe P_E_ |
| gvmE-pR | AGTTCCACGTCGCCGTACAGCT |  |
| gvmpF | AACATGGTCAAGCCGCTCCG | Probe P_F_ |
| gvmF-pR | ACGGCGTAACGCTCGCGTAT |  |
| 1133/34-pF | GCTTGTGACAGTGCCTTGAA | Probe P_1133-1134_ |
| 1133/34-pR | GGTGAGGATGGCGTATCG |  |
| 1543/44-pF | ATTGCAGACGCCGTAGACCG | Probe P_1544-1543_ |
| 1543/44-pR | TCGCCCAGGACGAGCGAGAC |  |
| 0026-pF | GCGTCGCGGAGTTCGAGGTGAA | Probe P_0026_ |
| 0026-pR | ACAGCGGCCCTGCCTCTGTT |  |
| 0174-pF | GGCCACTCCCGGACATCA | Probe P_0174_ |
| 0174-pR | GGCCGGTCGTTCTTGGAT |  |
| 0741-pF | GACCCCAGCAGTCACCACA | Probe P_0741_ |
| 0741-pR | CCACGACCCAGAAGAACCAG |  |
| 1209-pF | TCGCCGAGCATCTCCTGT | Probe P_1209_ |
| 1209-pR | GTTCATTGACCCGCACCC |  |
| 1410-pF | GATTTCCTTGACGGTGGCG | Probe P_1410_ |
| 1410-pR | GGACAGCAGGGCTTGTTCG |  |
| 1744-pF | ACCGCCTGCGCCGAAGTGAT | Probe P_1744_ |
| 1744-pR | CCCGTGCGTGGCTGGAAAT |  |
| 1780-pF | AACCCAGTTGCCGAGGACA | Probe P_1780_ |
| 1780-pR | TCCACAGCACGACGGTCAG |  |
| 1876-pF | GGTCGCCAGCACCAGCCA | Probe P_1876_ |
| 1876-pR | CTGGAAGAGAAGGTAGGCGGCC |  |
| 1920-pF | GGCAATCTTCCCGGCGCGC | Probe P_1920_ |
| 1920-pR | GGGTGCTCCTCAGCGATGCC |  |
| 1939-pF | GGCGACACCACCAACACCA | Probe P_1939_ |
| 1939-pR | ATGAGCTGGACAATTCCGTAGA |  |
| 1991-pF | CGATCAGCTCCACGCCCGGATA | Probe P_1991_ |
| 1991-pR | GGCGACGAAACGAGGCGC |  |
| 2100-pF | GAGCTGGCAGGTCAGAGAGCT | Probe P_2100_ |
| 2100-pR | GGCTCCAGGTGGCCAGCCA |  |
| 2106-pF | GGAAGCCCACCACGTTGACGAT | Probe P_2106_ |
| 2106-pR | ACCAGTTCGTCCGGGCGG |  |
| 2320-pF | TCACCGCATCCCGCCAGG | Probe P_2320_ |
| 2320-pR | CCGTCAGCTCGTGCTCCAC |  |
| 2888-pF | GCCCCACCCCTGCTGTGA | Probe P_2888_ |
| 2888-pR | TGCGGTCGAGTCGTCTGC |  |
| 2961-pF | AAGTGCTCGGTGGGAACATGC | Probe P_2961_ |
| 2961-pR | TCGGCGATGATCCAGCGGT |  |
| 3087-pF | TGGTGATCTCGATGAACTCTGG | Probe P_3087_ |
| 3087-pR | CCGTCCGAATCCTGGTGC |  |
| 3181-pF | ACAGGCCTACGTACCCACC | Probe P_3181_ |
| 3181-pR | GCACTTCGCAGTGCGGCA |  |
| 3187-pF | CACTCCCACCGTGACAAAGA | Probe P_3187_ |
| 3187-pR | GACCAGCGAAAAGGACAAAA |  |
| 3479-pF | ACTGGGTGATCTTCGCCTGGC | Probe P_3479_ |
| 3479-pR | CCAGGGCGAGGTCTCCGT |  |
| 3524-pF | GCGATGCCGAACGGGATG | Probe P_3524_ |
| 3524-pR | CGAAATCGGCACCCAGCA |  |
| 3535-pF | TAGCGGACCTGGGTGTGCG | Probe P_3535_ |
| 3535-pR | TGCGCAAGGCCACCGAGC |  |
| 3567-pF | AGCATCGCCGTCAGGTAGTACTT | Probe P_3567_ |
| 3567-pR | GGTGAGGAAGCCACCGGTCAC |  |
| 3624-pF | CTGATCGGTGGGGAAGTGGAGG | Probe P_3624_ |
| 3624-pR | ACCCGCGTGAAGAGACCGG |  |
| 3682-pF | ACCCGGAGTCCGATATGAGGC | Probe P_3682_ |
| 3682-pR | CGGTAGGCGAGAAGGAAGGAAA |  |
| 3696-pF | CACGGTGCTGCAGCGCAT | Probe P_3696_ |
| 3696-pR | CCCTGCTGAGCTGAGGCGAC |  |
| 3709-pF | GGACGTCGGTGGAGAGGACG | Probe P_3709_ |
| 3709-pR | TGGTCCCCCGCCCTCGAC |  |
| 3712-pF | CGTTGACCCCGTGCAATTC | Probe P_3712_ |
| 3712-pR | CCAGTCACCGAGCGGAAAG |  |
| 4030-pF | GCGGTGCCGTTCGGTGTG | Probe P_4030_ |
| 4030-pR | CACCAGAGCGGCGAGGGAGA |  |
| 4384-pF | GCACACCCGTCAGCAGCTC | Probe P_4384_ |
| 4384-pR | ACGCGACGGTGTAGGCGAT |  |
| 4507-pF | GTCGAGACGGCCGAGTTCCC | Probe P_4507_ |
| 4507-pR | GGCGATGGAGCCCACGGAA |  |
| 4595-pF | CGGGTCGGTGCTGTCTGC | Probe P_4595_ |
| 4595-pR | CGATCACCTCCAGGAGCG |  |
| 4691-pF | AGGCCACGGTGGCCGTCA | Probe P_4691_ |
| 4691-pR | CAGAAAGCCGCCCAGCAG |  |
| 4714-pF | CGGTCCGAGTCGGGGGAG | Probe P_4714_ |
| 4714-pR | GACGGCCACGATCTCGGC |  |
| 4782-pF | GTGCGCAGGGTGTGCTGCA | Probe P_4782_ |
| 4782-pR | GCACCGTCCGTACCCGGAT |  |
| 4948-pF | CTAACCGGTTCAGCCCCTCGTA | Probe P_4948_ |
| 4948-pR | TCCCGCAGCTCCCGGACG |  |
| 4997-pF | GATTTCGGCGGTGGTGCT | Probe P_4997_ |
| 4997-pR | GCGGGCTGATCGGTCATA |  |
| 5103-pF | AGCTCACGCTGCACCAGC | Probe P_5103_ |
| 5103-pR | CGCATTTCCGCCACCATC |  |
| 5223-pF | TGGGGCTGCTGCTGTACACG | Probe P_5223_ |
| 5223-pR | TTCTTGCCGAGCGCGTCGT |  |
| 5227-pF | CGCGTCGTGATCCGCCTTG | Probe P_5277_ |
| 5227-pR | CGATCTCGCGCTCTTCCAGGT |  |
| 5331-pF | ACCTGCTCGGTGACGGACTG | Probe P_5311_ |
| 5331-pR | GTTGACGATGCCCTTGGTG |  |
| 5381-pF | CCTGGAGGCCAGGGACGG | Probe P_5381_ |
| 5381-pR | CGAGGAGGGCTTGAGCTCGAT |  |
| 5700-pF | GGTCTGGGCGATCTGCAC | Probe P_5700_ |
| 5700-pR | GGTCTCCCCGAGCTTGTCA |  |
| 5778-pF | TCGTCGAGGACTTCACCAGC | Probe P_5778_ |
| 5778-pR | GAGAAACGTGGCGAGGGA |  |
| 5806-pF | GACGGCACGTCGGGTGCG | Probe P_5806_ |
| 5806-pR | TGGACGGCAGGTATGGCGG |  |
| 5926-pF | CTCGCCGCTGGGTATGTT | Probe P_5926_ |
| 5926-pR | TGCCCGCCAAGTGAAGTG |  |
| 6201-pF | ACGAGGTGGATGTCGGTGTC | Probe P_6201_ |
| 6201-pR | CGTGCCAGGTGCGGTAGAA |  |
| 6257-pF | CGGCGTCCATGTATCCGG | Probe P_6257_ |
| 6257-pR | GGAGTTGGGCCCGTCATC |  |
| 6438-pF | CCTGCCAGTGACGGGAGCCT | Probe P_6438_ |
| 6438-pR | GCCCAGCCGGTGGAACAA |  |
| 6724-pF | CTTCGTGGTCCCCGGAGAGC | Probe P_6724_ |
| 6724-pR | GCAGCAGCTGCTGGACCT |  |
| 6774-pF | GTCGGTGCCGTGTTCCAT | Probe P_6774_ |
| 6774-pR | TGAGCAGTGCCTCGTGCA |  |
| 6992-pF | CAGGACGCGCTCGTAATCGG | Probe P_6992_ |
| 6992-pR | TCCACGACACGAGGCACACG |  |
| 7577-pF | CATGGTGTTCGTGGTGAGC | Probe P_7577_ |
| 7577-pR | CGATCGGGACCAGCATCT |  |
| **For GUS assays** | | |
| gvmT1GUSA-pF | **TGCCAAGCTTGGGCTGCAGGTCGAC**AAGGCGGTGCTGATCTTCGTA | *gvmT1* promoter |
| gvmT1GUSA-pR | **TTTCGACGGGCCGCAGACCGGTCAT**CTCGTCACTCGGTTCCACTGGT |  |
| gvmT2GUSA-pF | **TTTCGACGGGCCGCAGACCGGTCAT**GAAGAAAGAAACTCCTTCGGCAGGG | *gvmT2* promoter |
| gvmT2GUSA -pR | **TGCCAAGCTTGGGCTGCAGGTCGAC**GGCTGGGGCACCGAGACATC |  |
| gvmEGUSA -pF | **TGCCAAGCTTGGGCTGCAGGTCGAC**AGGTGGGCCTGGTCATCGA | *gvmE* promoter |
| gvmEGUSA -pR | **TTTCGACGGGCCGCAGACCGGTCAT**GAGCAAGGGTCCTTGATCTTCGT |  |
| gvmFGUSA -pF | **TGCCAAGCTTGGGCTGCAGGTCGAC**CCTACGTGATCGCCCGCAAGAA | *gvmF* promoter |
| gvmFGUSA -pR | **TTTCGACGGGCCGCAGACCGGTCAT**CGTGCAACTCCTAGGGGTTCGTATG |  |
| gusA-F | ATGACCGGTCTGCGGCCCGTC | *gusA* ORF |
| gusA-R | **CAGGAAACAGCTATGACATGATTAC**TCACTGCTTCCCGCCCTGCTGC |  |
| NheI-F | GCGGACGCTAGCGCGGTTGCTCAACCAGAGATAACAGT | Conserved binding sequence confirmation |
| NheI-R | TACCAGATCATACGTATGATGTCCGGCG |  |
| NdeI-F | GCGGACATCATACGTATGATGCGGT |  |
| NdeI-R | TACCAGCATATGGTCCGGCGCAAGATCTGGTGA |  |
| PgvmA-XF | **CAGGCTGCGCAACTGTTGGGAAGGG** GGTGGCCTGTGACACGCCA |  |
| PgvmA-XR | **TTTCGACGGGCCGCAGACCGGTCAT**  TCCGGACCGGGCCAACGC |  |
| GusA-R | **CGATATCGCGCGCGGCCGCGGATCC**  TCACTGCTTCCCGCCCTGCTGC |  |
| PhrdB-F | **GCCGCAGCAGGGCGGGAAGCAGTGA**CCGCCTTCCGCCGGAACGG |  |
| PhrdB-R | GAACAACCTCTCGGAACGTTGAAAA |  |
| HPgvmR-F | **TTTTCAACGTTCCGAGAGGTTGTTC**TTGTCTGAAGGACTCCACAGGGCA |  |
| HPgvmR-R | **ACAGCTATGACATGATTACGAATTC**AGAGGCATGAAGAAAGAAACTCCTTCG |  |
| The introduced restriction sites are underlined. The overlapped sequences used for one step cloning are in bold. | | |

| **Table S2** Genes whose promoter regions contain sequences similar to GvmR binding sites | | | |
| --- | --- | --- | --- |
| # | Locus tag | Function | [Sequence](http://www.baidu.com/link?url=d-hL8dJ8uP7N68zv2rqvfvrSl9ai6pUsSAlpMm0PQAI6_6mETq9yJQDSdq2NEYQnzc_SV37owcgi_ssc4YOpHQhx_viKBI3a_mAxMbJ1Djr6tK_iR0RsVLoJrt2wQbeY) |
| 1 | *scn1544* | LacI family transcriptional regulator | gtgatacgtataacn_7_ gttatacgtatcat |
| 2 | *scn4948* | methyltransferase | atcctacgtttgaa |
| 3 | *scn5806* | KsbA | atcctccgtatgct |
| 4 | *scn0174* | major facilitator transporter | gccgtacgcgccac |
| 5 | *scn6257* | monooxygenase | ctcatacgtcggat |
| 6 | *scn3087* | hypothetical protein | gtcacgcggatgac |
| 7 | *scn4691* | hypothetical protein | atcatcggcatcgc |
| 8 | *scn0026* | hypothetical protein | gggatacggatgac |
| 9 | *scn1744* | hypothetical protein | gtcatacgcctcgc |
| 10 | *scn3187* | transglycosylase | gtcacacgactgac |
| 11 | *scn0741* | amino acid transporter | gtgatgcggctcac |
| 12 | *scn3682* | peptidase | gttgtccgcatcac |
| 13 | *scn5381* | branched-chain amino acid aminotransferase | gttcaacggatcgt |
| 14 | *scn5926* | hypothetical protein | ctgatgcgtgtgat |
| 15 | *scn5103* | membrane protein | gtgattcgactgac |
| 16 | *scn1939* | ABC transporter permease | gacatacgaacgac |
| 17 | *scn6438* | hypothetical protein, partial | gtgatgcgaaacat |
| 18 | *scn6201* | PucR family transcriptional regulator | aagatcctcatctc |
| 19 | *scn3624* | AraC family transcriptional regulator | atgtcttgcatgtc |
| 20 | *scn6724* | hypothetical protein, partial | gtccgccgcaccac |
| 21 | *scn3181* | NADH dehydrogenase | attgttcgtccaac |
| 22 | *scn1410* | preprotein translocase subunit YajC | atccttcgcacgac |
| 23 | *scn4997* | 1D-myo-inositol 2-acetamido-2-deoxy-alpha-D-glucopyranoside deacetylase | gctgtcggtatgac |
| 24 | *scn5331* | ABC transporter substrate-binding protein | catcatgn_32_catcacc |
| 25 | *scn7577* | hypothetical protein | gagatgcgtatgac |
| 26 | *scn5700* | DNA-binding protein | ggatgacn_20_gtcattc |
| 27 | *scn4595* | hypothetical protein | gcatcacn_10_gtcatcg |
| 28 | *scn2320* | hypothetical protein | gaatgacn_87_gtcatcc |
| 29 | *scn2888* | sugar ABC transporter substrate-binding protein | gtaccacn_49_gtatcac |
| 30 | *scn1920* | helicase | gagtgacn_3_atcactc |
| 31 | *scn5778* | PTS ascorbate transporter subunit IIC | gccatacn_163_gtacaac |
| 32 | *scn1780* | signal peptidase | gcatgacn_17_gtgatgc |
| 33 | *scn6774* | hypothetical protein | ctgattcn_60_gtcatgc |
| 34 | *scn3524* | membrane protein | atgatgcn_93_gcatcgg |
| 35 | *scn1209* | membrane protein | gtggtacn_7_gtggtac |
| 36 | *scn3567* | MULTISPECIES: NADH dehydrogenase subunit A | ggatgagn_534_ctcatac |
| 37 | *scn3535* | MULTISPECIES: NADH-ubiquinone oxidoreductase subunit 3 | ggatgggn_129_ggatcgg |
| 38 | *scn4384* | NADH-quinone oxidoreductase subunit D | gggatgcn_45_ccgatgc |
| 39 | *scn1991* | succinate dehydrogenase | gatgagcn_16_ctgatgg |
| 40 | *scn4782* | succinate dehydrogenase | gtgtgatn_20_gtgtgat |
| 41 | *scn2100* | cystathionine beta-lyase | ggattacn_67_ggactac |
| 42 | *scn2106* | cytochrome C oxidase subunit II | ccttatan_196_tttatgc |
| 43 | *scn1876* | protoheme IX farnesyltransferase | ggtcatcn_99_ctgatgt |
| 44 | *scn4030* | ABC transporter | tcatcacn_28_gtgatgc |
| 45 | *scn5227* | ATP synthase F0F1 subunit alpha | gatccggn_8_gatccgg |
| 46 | *scn5223* | ATP synthase F0F1 subunit A | ccgatgcn_107_cccatgc |
| 47 | *scn4507* | polyphosphate kinase | ctgatacn_16_ctgatac |
| 48 | *scn3712* | phosphoribosylamine--glycine ligase | gcatgcan_45_gtcatcc |
| 49 | *scn3696* | phosphoribosylaminoimidazole synthetase | atgacggn_229_atcatgc |
| 50 | *scn2961* | phosphoribosylaminoimidazole carboxylase | gtcatacn_91_ggatgat |
| 51 | *scn3709* | phosphoribosylaminoimidazole-succinocarboxamide synthase | ttgatccn_10_tcgattc |
| 52 | *scn6992* | adenylosuccinate lyase | gcatgagn_32_gcatgtg |
| 53 | *scn4714* | phosphoribosylaminoimidazolecarboxamide formyltransferase | gcatgagn_1_gcatcaa |
| 54 | *scn3479* | adenylate kinase, partial | gcatcagn_81_aggatgc |

The space length between each pair of inverted repeats is indicated by red color font.

**Table S3** Comparative transcriptional analysis of genes involved in oxidative phosphorylation

| Gene ID | log_2_fold change (0.5d) | log_2_fold change (1d) | log_2_fold change (2d) | log_2_fold change (5d) |
| --- | --- | --- | --- | --- |
| *scn3181* | 0.075186146 | 0.228577758 | -2.847550204 | -1.675784016 |
| *scn3526* | -0.352481344 | 1.252017791 | 0.125924529 | -1.016279773 |
| *scn3527* | -0.390698973 | 1.203461607 | 0.310435149 | -0.410417533 |
| *scn3528* | -0.41521506 | 1.217540906 | 0.110226903 | -0.988099927 |
| *scn3529* | -0.050638334 | 1.027177018 | 0.18403166 | -0.989895415 |
| *scn3530* | -0.576268453 | 0.997812511 | 0.127500175 | -0.478685212 |
| *scn3531* | -0.553024957 | 1.600989574 | -0.05213316 | -0.376609911 |
| *scn3532* | -0.570561268 | 1.399602692 | 0.098574917 | -0.862382149 |
| *scn3534* | -0.655764972 | 1.61354516 | 0.170051269 | -0.521919054 |
| *scn3554* | 1.499960794 | -6.034508991 | 0.853259234 | -1.075757996 |
| *scn3555* | 1.624895044 | -5.850443599 | 0.961298965 | -0.8239642 |
| *scn3556* | 1.660012901 | -5.550082632 | 0.973043188 | -0.902567918 |
| *scn3557* | 2.22310228 | -6.577081676 | 1.166306318 | -0.357958741 |
| *scn3558* | 2.048783693 | -5.536876335 | 1.11259816 | -0.815942631 |
| *scn3559* | 1.446853024 | -5.417637388 | 0.994218869 | -0.641437475 |
| *scn3560* | 1.377819023 | -6.34704512 | 1.106575351 | -0.540462348 |
| *scn3561* | 1.421686713 | -6.516048168 | 1.080829322 | -1.080257326 |
| *scn3562* | 0.976666482 | -6.29991004 | 1.121520171 | -0.643507783 |
| *scn3563* | 1.258415966 | -6.390631359 | 1.271676944 | -0.922025473 |
| *scn3564* | 1.458665801 | -6.245790014 | 1.19683472 | -0.681598283 |
| *scn3565* | 2.291119512 | -6.854650225 | 1.153982465 | -1.055841579 |
| *scn3566* | 1.831098188 | -6.695543795 | 1.358671334 | -0.707613078 |
| *scn3567* | 2.047157478 | -7.445528157 | 1.139272807 | -0.963287631 |
| *scn3664* | 0.537415921 | -1.662448451 | -1.774511819 | 0.095458929 |
| *scn4384* | -0.154659512 | 1.525414798 | -0.559875816 | -1.044818906 |
| *scn1991* | 0.737167505 | 0.54159192 | -2.732726318 | -2.882898593 |
| *scn1992* | 0.780205901 | 0.086437647 | -2.957574082 | -2.730166873 |
| *scn1993* | 0.850410993 | 0.177007125 | -2.641543374 | -2.427342563 |
| *scn4779* | -0.46118533 | 0.966294245 | 0.073557017 | -0.963235891 |
| *scn4780* | -0.575633026 | 1.044449459 | 0.04514443 | -1.090114458 |
| *scn4781* | -0.49556985 | 1.01633009 | -0.085741045 | -1.384381623 |
| *scn4782* | -0.582465514 | 0.945352118 | -0.129282732 | -1.354456365 |
| *scn4977* | -0.788795244 | -2.653099589 | -0.233157748 | 0.51892048 |
| *scn4978* | 0.393006499 | -2.670503662 | -0.242654025 | 0.036206332 |
| *scn4028* | 0.558402086 | 4.184377764 | -4.650848233 | -1.707669888 |
| *scn2098* | -0.445986677 | 0.506757699 | -0.370774304 | -1.407285299 |
| *scn2099* | -0.408967038 | 0.513782255 | -0.317440074 | -1.415542687 |
| *scn2100* | -0.413499862 | 0.56553744 | -0.463839155 | -1.521327475 |
| *scn1876* | -0.223418613 | 0.8293208 | -0.815017164 | -1.528892269 |
| *scn2101* | -0.359207665 | 0.507931849 | -0.676224671 | -1.7305902 |
| *scn2105* | -0.369209551 | 0.941189736 | -0.432207291 | -1.415052529 |
| *scn2106* | -0.40647436 | 0.984337258 | -0.365559028 | -1.472992653 |
| *scn5223* | -0.849644179 | -0.310047465 | 0.307049185 | 0.638030451 |
| *scn5224* | -0.59020427 | -0.498049051 | 0.23628081 | 0.825552933 |
| *scn5225* | -0.346638028 | -0.429395649 | 0.260912697 | 0.548690178 |
| *scn5227* | -0.500158976 | -0.173212416 | 0.234515517 | 0.415632665 |
| *scn5228* | -0.336463419 | 0.01680595 | 0.31548019 | 0.677827677 |
| *scn5229* | -0.464690194 | -0.192622041 | 0.3318087 | 0.588602449 |

**Table S4** Comparative transcriptional analysis of genes involved in ribosome

| Gene ID | log_2_fold change (0.5d) | log_2_fold change (1d) | log_2_fold change (2d) | log_2_fold change (5d) |
| --- | --- | --- | --- | --- |
| *scn3511* | -0.686693191 | 1.339772403 | 0.696049286 | -0.003528178 |
| *scn3496* | -0.923866688 | 1.210050571 | 0.6963646 | 0.279086778 |
| *scn3499* | -1.072201342 | 1.250723099 | 0.687487112 | 0.31584201 |
| *scn3498* | -0.849256795 | 1.226817418 | 0.686370753 | 0.44105932 |
| *scn3487* | -0.905601473 | 1.22874707 | 0.780618015 | 0.569886211 |
| *scn3485* | -0.716638255 | 1.296670618 | 0.673116782 | 0.630806565 |
| *scn3508* | -0.585265714 | 1.107637993 | 1.122395407 | 0.156605084 |
| *scn3994* | -0.651703218 | 1.240821207 | 0.824065393 | 0.820758104 |
| *scn3509* | -0.680870636 | 1.259171131 | 1.153333918 | 0.223942351 |
| *scn3512* | -0.781646788 | 1.078468049 | 0.51334127 | 0.29876456 |
| *scn3467* | -0.831580107 | 0.898438887 | 0.624486709 | 0.123615986 |
| *scn3489* | -0.976502288 | 1.288018936 | 0.8058115 | 0.307112961 |
| *scn3481* | -0.773045133 | 1.205295591 | 0.531780512 | 0.502848949 |
| *scn3492* | -0.924871205 | 1.232436083 | 0.820787137 | 0.329759869 |
| *scn3471* | -0.575145451 | 1.345477207 | 0.694737011 | 0.898480352 |
| *scn3484* | -0.711199507 | 1.312408559 | 0.59182076 | 0.272674057 |
| *scn5465* | -0.606489339 | 0.901349998 | 0.418889767 | 0.402644116 |
| *scn1504* | -0.595608328 | 0.840924593 | 0.892990948 | 0.989336541 |
| *scn3494* | -0.996887722 | 1.217412192 | 0.726285632 | 0.416725147 |
| *scn3497* | -1.003408565 | 1.256766689 | 0.701768149 | 0.718142954 |
| *scn3488* | -0.97806348 | 1.256499292 | 0.879166531 | 0.696932637 |
| *scn2541* | -0.614739795 | 0.902179133 | 0.625781502 | 0.865001401 |
| *scn5424* | -0.743538935 | 0.635672334 | 0.58947026 | 0.941329437 |
| *scn3491* | -0.905277347 | 1.21815877 | 0.779332251 | 0.454362157 |
| *scn3482* | -0.844057037 | 1.298834248 | 0.47125417 | 0.010540068 |
| *scn4182* | -0.17344761 | -0.013335325 | 3.143416012 | 1.611508899 |
| *scn5431* | -0.631184546 | 0.854093486 | 0.879099035 | 0.816189006 |
| *scn3520* | -0.748026344 | 1.708087734 | 0.635350811 | 0.937837088 |
| *scn4346* | 0.893993326 | -1.501211375 | 2.009973483 | 1.441158144 |
| *scn3965* | -0.755893766 | 1.28935253 | 1.152942738 | 0.169318997 |
| *scn1505* | -0.585995257 | 0.941091833 | 0.736021974 | 0.878111893 |
| *scn3476* | -0.553980543 | 0.939977999 | 0.548235764 | 0.434332948 |
| *scn1927* | -0.683377085 | 0.587660545 | 0.615588437 | 0.086754621 |
| *scn5477* | -0.727368451 | 1.090909454 | 0.697170303 | 0.200456526 |
| *scn3493* | -0.812119712 | 1.220763452 | 0.767610297 | 0.318987518 |
| *scn1397* | -0.491718782 | 1.334796744 | 0.241380705 | -0.638177171 |
| *scn3483* | -0.697050762 | 1.255087568 | 0.601347615 | 0.414186318 |
| *scn3991* | -0.714031796 | 1.112284808 | 0.829502274 | 0.338898182 |
| *scn3503* | -0.793616547 | 0.972610779 | 0.742124961 | 0.556836451 |
| *scn3486* | -0.812525189 | 1.247534388 | 0.539074616 | 0.30403463 |
| *scn3466* | -0.788429771 | 0.961146291 | 0.678315136 | 0.426544628 |
| *scn3500* | -0.989073512 | 1.205700255 | 0.52319719 | 0.407538147 |
| *scn3474* | -0.666774898 | 0.90266789 | 0.508788908 | 0.547349925 |
| *scn3504* | -0.795828752 | 1.033658978 | 0.866621075 | 0.680846277 |
| *scn3475* | -0.724621328 | 0.852440553 | 0.509805577 | 0.749354372 |
| *scn5551* | -0.579383989 | 1.051769647 | 0.755299705 | 0.632186784 |
| *scn5461* | -0.737204748 | 0.828153593 | 0.51456251 | 0.046765411 |
| *scn3490* | -0.811246742 | 1.27749078 | 0.837872384 | 0.723326804 |
| *scn3993* | -0.603015456 | 1.177762684 | 0.737863085 | 0.579751599 |
| *scn3495* | -0.987838926 | 1.184951318 | 0.689533438 | 0.606189119 |

**Table S5** Comparative transcriptional analysis of genes involved in purine biosynthesis

| Gene ID | log_2_fold change (0.5d) | log_2_fold change (1d) | log_2_fold change (2d) | log_2_fold change (5d) |
| --- | --- | --- | --- | --- |
| *scn3207* | -0.384268012 | 0.960965419 | -0.505461355 | -0.088570616 |
| *scn3697* | -0.527946184 | 1.006557506 | -0.718146434 | -1.615378292 |
| *scn3712* | 0.111944149 | 1.8427425 | -0.418779208 | -0.685814543 |
| *scn3696* | -0.505591774 | 1.14880715 | -0.457660075 | -1.508190265 |
| *scn2961* | -0.508226132 | 0.696250203 | -0.527282676 | -1.166389814 |
| *scn2960* | -0.277340387 | 0.494139386 | -0.236529273 | -1.09242725 |
| *scn3709* | -0.073173408 | -0.412521817 | -0.118526698 | -0.56539848 |
| *scn6992* | -0.36097276 | 0.772150424 | -0.60493823 | -1.135785103 |
| *scn4714* | -0.277235942 | 0.526547825 | 0.011140262 | -0.89006062 |
| *scn3722* | -0.448223477 | 1.050406771 | 0.212856888 | -0.175983197 |
| *scn1359* | -0.24686065 | 2.774221605 | -0.702930111 | -0.506817963 |
| *scn3436* | -0.249342882 | 1.186277205 | 0.324340803 | -0.01716771 |
| *scn4673* | -0.48472824 | 0.662799371 | -0.127549548 | 0.225286462 |

**Table S6** Comparative transcriptional analysis of 49 transcriptional regulatory genes

| Gene ID | Type of product | log_2_fold change (0.5d) | log_2_fold change (1d) | log_2_fold change (2d) | log_2_fold change (5d) |
| --- | --- | --- | --- | --- | --- |
| *scn0235* | DeoR/GlpR family | 0.125849256 | 2.134422399 | 0.048936551 | 0.043657887 |
| *scn0266* | MarR family | 0.409454666 | 3.17255763 | 0.236698037 | 0.632476887 |
| *scn0505* | NarL/FixJ family | 0.975954468 | -2.693635596 | -0.212862332 | -0.153771618 |
| *scn0872* | OmpR family | -1.00505263 | -3.67178409 | 0.722384924 | -0.248100368 |
| *scn1024* | IclR family | -0.284122289 | -0.666579261 | -0.72703966 | 2.493476042 |
| *scn1357* | Lrp family | -0.592208267 | 2.62255733 | 0.859588569 | -0.547517475 |
| *scn1561* | IclR family | 0.343798241 | 2.838724066 | 0.760278683 | 0.511200837 |
| *scn1645* | GntR family | 0.945022085 | -2.68342146 | -0.433315106 | -0.094715608 |
| *scn1702* | YafY family | 0.040004366 | 1.032970101 | -0.803018908 | -2.4555881 |
| *scn1913* | MolR family | 1.928204879 | -0.423761813 | 3.026108712 | 1.343301892 |
| *scn2349* | AcrR family | -0.366826186 | 3.205293952 | 1.112548677 | 0.877805627 |
| *scn2360* | PucR family | -0.008993318 | 1.609178478 | -0.705709389 | -2.024837607 |
| *scn2470* | XRE-family | 0.696531865 | -4.226336693 | -0.123702166 | 1.705582321 |
| *scn2688* | AcrR family | -0.627947805 | 6.441377766 | 1.432476586 | -0.398002173 |
| *scn2833* | Lrp family | 0.634151219 | -2.306030907 | -0.635198934 | -0.507214999 |
| *scn2903* | AcrR family | -0.330243662 | 2.029759551 | 1.095717915 | 0.121778194 |
| *scn3220* | NarL/FixJ family | 1.093859971 | -2.863796451 | 0.034948035 | 0.5489582 |
| *scn3226* | NarL/FixJ family | 0.397242768 | -3.072458347 | 0.716559803 | -0.414708793 |
| *scn3254* | AcrR family | -0.499665764 | 3.093952132 | 0.516503724 | 0.702238729 |
| *scn3328* | DeoR/GlpR family | 1.146083115 | 8.487996751 | 2.892604107 | -0.330426434 |
| *scn3340* | AcrR family | 0.069827497 | 5.944870732 | -2.749076865 | -1.685286292 |
| *scn3360* | XRE-family | -2.734669269 | -8.309159587 | -9.643315192 | -7.114455953 |
| *scn3439* | NarL/FixJ family | 1.839658368 | -5.829899069 | 0.21440269 | 1.078501834 |
| *scn3655* | AlpA family | 0.884325578 | -3.361905788 | 0.744507508 | 0.256987817 |
| *scn3811* | NarL/FixJ family | 0.23068921 | -2.401842091 | 0.643683794 | 0.039940669 |
| *scn3915* | AcrR family | -0.633713125 | -2.539948131 | 0.213555996 | 1.02819755 |
| *scn4050* | FadR family | -0.566270535 | 2.120424598 | 0.227275435 | -0.38500915 |
| *scn4156* | MarR family | -0.650380313 | 4.494773107 | 0.389051184 | -1.537032208 |
| *scn4352* | XRE-family | -0.34705214 | -2.354831817 | 0.695742621 | 0.838451659 |
| *scn4452* | FrmR family | -0.267021136 | 0.600256134 | -0.882903976 | 2.04204532 |
| *scn4493* | OmpR family | 0.721371577 | 0.033155108 | -2.643935894 | -0.977452503 |
| *scn4586* | XRE-family | 1.269957748 | -4.225447037 | -0.979288902 | 1.209171422 |
| *scn4836* | DeoR/GlpR family | 1.379197377 | -4.535297241 | 0.208794889 | 1.063124223 |
| *scn4952* | XRE-family | 0.497949314 | -4.318897155 | 0.608332076 | 1.014345215 |
| *scn4970* | GntR family | 1.792533043 | -4.709898739 | 0.339909819 | 0.416232627 |
| *scn5162* | PadR family | -1.090239031 | 2.395717215 | 0.362587969 | -0.290582669 |
| *scn5207* | PleD family | -1.915621938 | -4.151477949 | -4.197802757 | -3.82518508 |
| *scn5264* | ArsR family | 0.186421901 | 0.089751475 | -2.187302728 | -3.076279212 |
| *scn5277* | AcrR family | -0.732448373 | 2.79629323 | -1.348090963 | -0.864510586 |
| *scn5806* | AcrR family | -1.144999218 | -3.865822359 | -0.484143971 | -0.247404238 |
| *scn5856* | NarL/FixJ family | 1.032500422 | -0.525821554 | -2.805220389 | 0.38227001 |
| *scn6001* | MarR family | -0.002334746 | 2.028672173 | -0.2146987 | -0.741286565 |
| *scn6296* | LysR family | 0.250037405 | -1.945381867 | -2.190511036 | 0.585800775 |
| *scn6328* | MmcQ/YjbR family | 1.794496468 | -0.170680803 | 2.019140784 | 0.475159134 |
| *scn6388* | MarR family | 2.18380302 | -7.840545183 | 0.475553865 | 1.572699978 |
| *scn6866* | AcoR | 1.431151799 | -3.182842058 | -1.674989663 | -0.300814984 |
| *scn7142* | NarL/FixJ family | 1.691132567 | -4.337836916 | -2.062156377 | -0.761115802 |
| *scn7547* | OmpR family | 0.62885155 | 0.925981258 | 3.052478403 | -0.672088386 |
| *scn7773* | MarR family | -0.563094009 | -2.466076637 | 0.747122555 | 0.282920246 |

**Table S7** The number of orthologs of the newly discovered guvermectin regulators in 268 *Streptomyces* genomes (as of Aug. 2021)

| Gene | Number of orthologs |
| --- | --- |
| *scn3360* | 263 |
| *scn6388* | 214 |
| *scn4970* | 252 |
| *scn4952* | 23 |
| *scn6293* | 154 |
| *scn4836* | 260 |

**Table S8** Strains and plasmids used in this study

| Strains/plasmids | Description | Source or reference |
| --- | --- | --- |
| *E. coli* | | |
| JM109 | General cloning host for plasmid manipulation | Novagen |
| ET12567/pUZ8002 | Non-methylating ET12567 containing non-transmissible RP4 derivative plasmid pUZ8002 | (1) |
| BL21(DE3) | Host for protein expression | Novagen |
| *Streptomyces* strains | | |
| *S. caniferus* | | |
| NEAU6 | Parental strain for guvermectin production | (2) |
| NEAU6/pIJ10500 | NEAU6 containing pIJ10500, Hyg^R^ | This work |
| ΔgvmR | *gvmR* inactivation strain | (2) |
| ΔgvmR/gvmR | ΔgvmR containing pIJ10500::gvmR | This work |
| ΔgvmR/P_hrdB_gvmR | ΔgvmR containing pIJ10500::P_hrdB_gvmR | This work |
| ΔgvmR/P_A_gusA | ΔgvmR containing pSyg::P_A_gusA | This work |
| ΔgvmR/P_A_gusA::P_hrdB_gvmR | ΔgvmR containing pSyg::P_A_gusA::P_hrdB_gvmR | This work |
| ΔgvmR/P_A_-M1gusA::P_hrdB_gvmR | ΔgvmR containing pSyg::P_A_-M1gusA::P_hrdB_gvmR | This work |
| ΔgvmR/P_A_-M2gusA::P_hrdB_gvmR | ΔgvmR containing pSyg::P_A_-M2gusA::P_hrdB_gvmR | This work |
| ΔgvmR/P_A_-M3gusA::P_hrdB_gvmR | ΔgvmR containing pSyg::P_A_-M3gusA::P_hrdB_gvmR | This work |
| OscnR1 | NEAU6 containing pSET152::P_hrdB_gvmR | This work |
| NEAU6/pSET152 | NEAU6 containing pSET152 |  |
| Oscn0235 | NEAU6 containing pSET152::P_hrdB_scn0235 | This work |
| Oscn0266 | NEAU6 containing pSET152::P_hrdB_scn0266 | This work |
| Oscn0505 | NEAU6 containing pSET152:: scn0505 | This work |
| Oscn0872 | NEAU6 containing pSET152::P_hrdB_scn0872 | This work |
| Oscn1024 | NEAU6 containingpSET152::P_hrdB_scn1024 | This work |
| Oscn1357 | NEAU6 containing pSET152::P_hrdB_scn1357 | This work |
| Oscn1561 | NEAU6 containing pSET152::P_hrdB_scn1561 | This work |
| Oscn1645 | NEAU6 containing pSET152::P_hrdB_scn1645 | This work |
| Oscn1702 | NEAU6 containing pSET152::P_hrdB_scn1702 | This work |
| Oscn1913 | NEAU6 containing pSET152::P_hrdB_scn1913 | This work |
| Oscn2349 | NEAU6 containing pSET152::P_hrdB_scn2349 | This work |
| Oscn2360 | NEAU6 containing pSET152::P_hrdB_scn2360 | This work |
| Oscn2470 | NEAU6 containing pSET152::P_hrdB_scn2470 | This work |
| Oscn2688 | NEAU6 containing pSET152::P_hrdB_scn2688 | This work |
| Oscn2833 | NEAU6 containing pSET152::P_hrdB_scn2833 | This work |
| Oscn2903 | NEAU6 containing pSET152::P_hrdB_scn2903 | This work |
| Oscn3220 | NEAU6 containing pSET152::P_hrdB_scn3220 | This work |
| Oscn3226 | NEAU6 containing pSET152::P_hrdB_scn3226 | This work |
| Oscn3254 | NEAU6 containing pSET152::P_hrdB_scn3254 | This work |
| Oscn3328 | NEAU6 containing pSET152::P_hrdB_scn3328 | This work |
| Oscn3340 | NEAU6 containing pSET152::P_hrdB_scn3340 | This work |
| Oscn3360 | NEAU6 containing pSET152::scn3360 | This work |
| Oscn3439 | NEAU6 containing pSET152::scn3439 | This work |
| Oscn3655 | NEAU6 containing pSET152::scn3655 | This work |
| Oscn3811 | NEAU6 containing pSET152::P_hrdB_scn3811 | This work |
| Oscn3915 | NEAU6 containing pSET152::P_hrdB_scn3915 | This work |
| Oscn4050 | NEAU6 containing pSET152::P_hrdB_scn4050 | This work |
| Oscn4156 | NEAU6 containing pSET152::P_hrdB_scn4156 | This work |
| Oscn4352 | NEAU6 containing pSET152::P_hrdB_scn4352 | This work |
| Oscn4452 | NEAU6 containing pSET152::P_hrdB_scn4452 | This work |
| Oscn4493 | NEAU6 containing pSET152::P_hrdB_scn4493 | This work |
| Oscn4586 | NEAU6 containing pSET152::P_hrdB_scn4586 | This work |
| Oscn4836 | NEAU6 containing pSET152::scn4836 | This work |
| Oscn4952 | NEAU6 containing pSET152::P_hrdB_scn4952 | This work |
| Oscn4970 | NEAU6 containing pSET152::P_hrdB_scn4970 | This work |
| Oscn5162 | NEAU6 containing pSET152::P_hrdB_scn5162 | This work |
| Oscn5207 | NEAU6 containing pSET152::P_hrdB_scn5207 | This work |
| Oscn5264 | NEAU6 containing pSET152::P_hrdB_scn5264 | This work |
| Oscn5277 | NEAU6 containing pSET152::P_hrdB_scn5277 | This work |
| Oscn5806 | NEAU6 containing pSET152::scn5806 | This work |
| Oscn5856 | NEAU6 containing pSET152::P_hrdB_scn5856 | This work |
| Oscn6001 | NEAU6 containing pSET152::P_hrdB_scn6001 | This work |
| Oscn6296 | NEAU6 containing pSET152::P_hrdB_scn6296 | This work |
| Oscn6328 | NEAU6 containing pSET152::P_hrdB_scn6328 | This work |
| Oscn6388 | NEAU6 containing pSET152::P_hrdB_scn6388 | This work |
| Oscn6866 | NEAU6 containing pSET152::P_hrdB_scn6866 | This work |
| Oscn7142 | NEAU6 containing pSET152::scn7142 | This work |
| Oscn7547 | NEAU6 containing pSET152::P_hrdB_scn7547 | This work |
| Oscn7773 | NEAU6 containing pSET152::P_hrdB_scn7773 | This work |
| M1146/P_T2_pRGUS | M1146 carrying the vector pT2GUS | This work |
| *S. coelicolor* |  |  |
| M1146 | Δact Δred Δcpk Δcda | (3) |
| M1146/gusA |  | This work |
| M1146/P_T1_gusA | M1146 containing pSET152::P_T1_gusA | This work |
| M1146/P_T2_gusA | M1146 containing pSET152::P_T2_gusA | This work |
| M1146/P_E_gusA | M1146 containing pSET152::P_E_gusA | This work |
| M1146/P_F_gusA | M1146 containing pSET152::P_F_gusA | This work |
| Plasmids |  |  |
| pIJ10500 | Integrative *E. coli*-*Streptomyces* shuttle vector | (1) |
| pSET152 | Integrative *E. coli*-*Streptomyces* shuttle vector | (1) |
| pBluescript II(KS+) | Routine cloning and subcloning vector | Novagen |
| pGEX4T-1 | Vector for GST-tagged protein expression in *E. coli* | GE Healthcare |
| pIJ10500:: gvmR | pIJ10500 containing one copy of *gvmR* driven by  its own promoter | This work |
| pIJ10500:: P_hrdB_gvmR | pIJ10500 containing one copy of *gvmR* driven by  the *hrdB* promoter | This work |
| pSyg::P_A_gusA | pSET152::hyg containing the coding region of *gusA* driven by the *gvmA* promoter | This work |
| pSyg::P_A_gusA::P_hrdB_gvmR | pSET152::hyg containing the coding region of *gusA* driven by P_A_ and one copy of *gvmR* driven by the *hrdB* promoter | This work |
| pSyg::P_A_-M1gusA::P_hrdB_gvmR | pSET152::hyg containing the coding region of *gusA* driven by P_A_-M1 and one copy of *gvmR* driven by the *hrdB* promoter | This work |
| pSyg::P_A_-M2gusA::P_hrdB_gvmR | pSET152::hyg containing the coding region of *gusA* driven by P_A_-M2 and one copy of *gvmR* driven by the *hrdB* promoter | This work |
| pSyg::P_A_-M3gusA::P_hrdB_gvmR | pSET152::hyg containing the coding region of *gusA* driven by P_A_-M3 and one copy of *gvmR* driven by the *hrdB* promoter | This work |
| pSET152::P_hrdB_scnR1 | pSET152 containing one copy of *scnR1* driven by  *hrdB* promoter | This work |
| pSET152::P_hrdB_scn0235 | pSET152 containing one copy of *scn0235* driven by *hrdB* promoter | This work |
| pSET152::P_hrdB_scn0266 | pSET152 containing one copy of *scn0266* driven by *hrdB* promoter | This work |
| pSET152::scn0505 | pSET152 containing one copy of *scn0505* driven by *hrdB* promoter | This work |
| pSET152::P_hrdB_scn0872 | pSET152 containing one copy of *scn0872* driven by *hrdB* promoter | This work |
| pSET152::P_hrdB_scn1024 | pSET152 containing one copy of *scn1024* driven by *hrdB* promoter | This work |
| pSET152::P_hrdB_scn1357 | pSET152 containing one copy of *scn1357* driven by *hrdB* promoter | This work |
| pSET152::P_hrdB_scn1561 | pSET152 containing one copy of *scn1561* driven by *hrdB* promoter | This work |
| pSET152::P_hrdB_scn1645 | pSET152 containing one copy of *scn1645* driven by *hrdB* promoter | This work |
| pSET152::P_hrdB_scn1702 | pSET152 containing one copy of *scn1702* driven by *hrdB* promoter | This work |
| pSET152::P_hrdB_scn1913 | pSET152 containing one copy of *scn1913* driven by *hrdB* promoter | This work |
| pSET152::P_hrdB_scn2349 | pSET152 containing one copy of *scn2349* driven by *hrdB* promoter | This work |
| pSET152::P_hrdB_scn2360 | pSET152 containing one copy of *scn2360* driven by *hrdB* promoter | This work |
| pSET152::P_hrdB_scn2470 | pSET152 containing one copy of *scn2470* driven by *hrdB* promoter | This work |
| pSET152::P_hrdB_scn2688 | pSET152 containing one copy of *scn2688* driven by *hrdB* promoter | This work |
| pSET152::P_hrdB_scn2833 | pSET152 containing one copy of *scn2833* driven by *hrdB* promoter | This work |
| pSET152::P_hrdB_scn2903 | pSET152 containing one copy of *scn2903* driven by *hrdB* promoter | This work |
| pSET152::P_hrdB_scn3220 | pSET152 containing one copy of *scn3220* driven by *hrdB* promoter | This work |
| pSET152::P_hrdB_scn3226 | pSET152 containing one copy of *scn3226* driven by *hrdB* promoter | This work |
| pSET152::P_hrdB_scn3254 | pSET152 containing one copy of *scn3254* driven by *hrdB* promoter | This work |
| pSET152::P_hrdB_scn3328 | pSET152 containing one copy of *scn3328* driven by *hrdB* promoter | This work |
| pSET152::P_hrdB_scn3340 | pSET152 containing one copy of *scn3340* driven by *hrdB* promoter | This work |
| pSET152::scn3360 | pSET152 containing one copy of *scn3360* driven by its own promoter | This work |
| pSET152::scn3439 | pSET152 containing one copy of *scn3439* driven by its own promoter | This work |
| pSET152::scn3655 | pSET152 containing one copy of *scn3655* driven by its own promoter | This work |
| pSET152::P_hrdB_scn3811 | pSET152 containing one copy of *scn3811* driven by *hrdB* promoter | This work |
| pSET152::P_hrdB_scn3915 | pSET152 containing one copy of *scn3915* driven by *hrdB* promoter | This work |
| pSET152::P_hrdB_scn4050 | pSET152 containing one copy of *scn4050* driven by *hrdB* promoter | This work |
| pSET152::P_hrdB_scn4156 | pSET152 containing one copy of *scn4156* driven by *hrdB* promoter | This work |
| pSET152::P_hrdB_scn4352 | pSET152 containing one copy of *scn4352* driven by *hrdB* promoter | This work |
| pSET152::P_hrdB_scn4452 | pSET152 containing one copy of *scn4452* driven by *hrdB* promoter | This work |
| pSET152::P_hrdB_scn4493 | pSET152 containing one copy of *scn4493* driven by *hrdB* promoter | This work |
| pSET152::P_hrdB_scn4586 | pSET152 containing one copy of *scn4586* driven by *hrdB* promoter | This work |
| pSET152::scn4836 | pSET152 containing one copy of *scn4836* driven by its own promoter | This work |
| pSET152::P_hrdB_scn4952 | pSET152 containing one copy of *scn4952* driven by *hrdB* promoter | This work |
| pSET152::P_hrdB_scn4970 | pSET152 containing one copy of *scn4970* driven by *hrdB* promoter | This work |
| pSET152::P_hrdB_scn5162 | pSET152 containing one copy of *scn5162* driven by *hrdB* promoter | This work |
| pSET152::P_hrdB_scn5207 | pSET152 containing one copy of *scn5207* driven by *hrdB* promoter | This work |
| pSET152::P_hrdB_scn5264 | pSET152 containing one copy of *scn5264* driven by *hrdB* promoter | This work |
| pSET152::P_hrdB_scn5277 | pSET152 containing one copy of *scn5277* driven by *hrdB* promoter | This work |
| pSET152::scn5806 | pSET152 containing one copy of *scn5806* driven by its own promoter | This work |
| pSET152::P_hrdB_scn5856 | pSET152 containing one copy of *scn5856* driven by *hrdB* promoter | This work |
| pSET152::P_hrdB_scn6001 | pSET152 containing one copy of *scn6001* driven by *hrdB* promoter | This work |
| pSET152::P_hrdB_scn6296 | pSET152 containing one copy of *scn6296* driven by *hrdB* promoter | This work |
| pSET152::P_hrdB_scn6328 | pSET152 containing one copy of *scn6328* driven by *hrdB* promoter | This work |
| pSET152::P_hrdB_scn6388 | pSET152 containing one copy of *scn6388* driven by *hrdB* promoter | This work |
| pSET152::P_hrdB_scn6866 | pSET152 containing one copy of *scn6866* driven by *hrdB* promoter | This work |
| pSET152::scn7142 | pSET152 containing one copy of *scn7142* driven by its own promoter | This work |
| pSET152::P_hrdB_scn7547 | pSET152 containing one copy of *scn7547* driven by *hrdB* promoter | This work |
| pSET152::P_hrdB_scn7773 | pSET152 containing one copy of *scn7773* driven by *hrdB* promoter | This work |
| pSET152::gusA | pSET152 containing the coding region of *gusA* | Laboratory stock |
| pSET152::P_T1_gusA | pSET152 containing the coding region of *gusA* driven by *gvmT1* promoter | This work |
| pSET152::P_T2_gusA | pSET152 containing the coding region of *gusA* driven by *gvmT2* promoter | This work |
| pSET152::P_E_gusA | pSET152 containing the coding region of *gusA* driven by *gvmE* promoter | This work |
| pSET152::P_F_gusA | pSET152 containing the coding region of *gusA* driven by *gvmF* promoter | This work |
| pBlu::P_R-A_ | pBluescript II(KS+) containing P_R-A_ | This work |
| pBlu::P_R-A_-M1 | pBluescript II(KS+) containing P_R-A_-M1 | This work |
| pBlu::P_R-A_-M2 | pBluescript II(KS+) containing P_R-A_-M2 | This work |
| pBlu::P_R-A_-M3 | pBluescript II(KS+) containing P_R-A_-M3 | This work |
| pGEX-4T-1::gvmR | GvmR expression vector based on pGEX-4T-1 | This work |
| pGEX-4T-1::scnR1 | ScnR1 expression vector based on pGEX-4T-1 | This work |
